# Elucidating the diversity of microeukaryotes and epi-endophytes in the brown algal holobiome

**DOI:** 10.1101/2021.05.09.443287

**Authors:** Marit F. Markussen Bjorbækmo, Juliet Brodie, Anders K. Krabberød, Ramiro Logares, Stephanie Attwood, Stein Fredriksen, Janina Fuss, Anders Wold-Dobbe, Kamran Shalchian-Tabrizi, David Bass

## Abstract

**Background:** Brown algae (Phaeophyceae) are essential species in coastal ecosystems where they form kelp forests and seaweed beds that support a wide diversity of marine life. Host-associated microbial communities are an integral part of phaeophyte biology. The bacterial microbial partners of brown algae have received far more attention than microbial eukaryotes. To our knowledge, this is the first study to investigate brown algal-associated eukaryotes (the eukaryome) using broadly targeting ‘pan-eukaryotic’ primers and high throughput sequencing (HTS). Using this approach, we aimed to unveil the eukaryome of seven large common brown algal species. We also aimed to assess whether these macroalgae harbour novel eukaryotic diversity and to ascribe putative functional roles to the host-associated eukaryome, based on taxonomic affiliation and phylogenetic placement.

**Results:** Our sequence dataset was dominated by brown algal reads, from the host species and potential symbionts. We also detected a broad taxonomic diversity of eukaryotes in the brown algal holobiomes, with OTUs taxonomically assigned to ten of the eukaryotic major Kingdoms or supergroups. A total of 265 microeukaryotic and epi-endophytic operational taxonomic units (OTUs) were defined, using 97% similarity cut off during clustering, and were dominated by OTUs assigned to stramenopiles, Alveolata and Fungi. Almost one third of the OTUs we detected have not been found in previous molecular environmental surveys, and represented potential novel eukaryotic diversity. This potential novel diversity was particularly prominent in phylogenetic groups comprising heterotrophic and parasitic organisms, such as labyrinthulids and oomycetes, Cercozoa, and Amoebozoa.

**Conclusions:** Our findings provide important baseline data for future studies of seaweed-associated microorganisms, and demonstrate that microeukaryotes and epi-endophytic eukaryotes should be considered as an integral part of brown algal holobionts. The potential novel eukaryotic diversity we found and the fact that the vast majority of macroalgae in marine habitats remain unexplored, demonstrates that brown algae and other seaweeds are potentially rich sources for a large and hidden diversity of novel microeukaryotes and epi-endophytes.

## Background

Seaweeds are essential to the health of our planet, providing core ecosystem services, and food and shelter to a wide diversity of marine life, from microbes to mammals, [1, 2 and references therein]. Brown algal seaweeds (Phaeophyceae) belong to the stramenopiles, a radiation of eukaryotes which is phylogenetically separated from red- and green algal seaweeds (Archaeplastida; Chlorophyta and Rhodophyta). The brown algae are one of the most diversified groups of benthic algae and comprise approximately 2000 species, encompassing complex multicellular species which have diversified and evolved since the Mesozoic Era ∼250 MYA [2, 3]. Large brown algae, especially those in the orders Laminariales, Fucales and Tilopteridales are dominant members of intertidal and shallow subtidal ecosystems worldwide, where they function as ecosystem engineers forming complex underwater forests. These coastal ecosystems are some of the most productive habitats on the planet [4], where seaweed beds and kelp forests are indispensable biodiversity hotspots, which also provide habitat and nursery for a wide range of taxa, including many commercially important species [5]. Brown algae have been shown to accommodate up to 100.000 individuals of different invertebrate species per square metre [6].

Brown algae and other seaweeds also harbour a wide diversity of microbial epibionts and endobionts, comprising eukaryotes [7–9], prokaryotes [10–12] and viruses [13–16]. These, together with the host, are collectively referred to as the seaweed holobiont and can be considered as a localized ecosystem living on and in a host [10, 17–19]. It is increasingly recognized that host-associated microbial communities (microbiomes) are an integral part of the host biology, exerting diverse and strong influences on their hosts [19, 20]. While the term microbiome usually refers mostly or exclusively to bacteria, it is important to consider the whole microbial symbiome, including microbial eukaryotes (microeukaryotes) and larger epi-endophytic symbionts, to enable a comprehensive understanding of holobiont functioning [19, 21].

Symbiotic interactions in its broadest sense refers to all intimate ecological interactions between two species [22]. According to the fitness effect on the members involved in the symbiotic relationship, the associations cover a wide spectrum of interactions along the gradual continuum between positive mutualism and negative parasitism. Some symbiotic interactions are obligate while other are more temporary and fluctuating, and the nature of the interaction can vary depending on the symbiont’s interactions with its hosts, other symbionts and environmental conditions [20]. To date, bacterial symbionts of brown algae have received most study [e.g., 10, 11, 12, 23], and have been shown to represent complex and highly dynamic relationships that can have everything from fundamental to detrimental effects on their hosts [10, 24–28].

Microeukaryotes in the brown algal symbiome are far less well known. Only a small number of studies have investigated these associations, mostly using traditional culturing/cell isolation methods or targeted molecular approaches. These studies demonstrate that a broad taxonomic diversity of microeukaryotes are associated with brown algae, including surface dwelling heterotrophic diatoms, dinoflagellates and ciliates [8], ‘naked amoeba’ [29], epiphytic and endophytic diatoms [30–32], and green algal endophytes [33], in addition to parasitic or saprotrophic labyrinthulids [34, 35], oomycetes [7, 36, 37], phytomyxids [38–40] and fungi [9, 41–44]. The nature of these microeukaryote-host relationships is mostly unknown, although some symbionts can have detrimental effects on their macroalgal hosts, for example phytomyxids [38–40], oomycetes [7, 36, 37] and chytridiomycete fungi [41]. Other microeukaryotes are suspected to have a beneficial effect on their hosts, for example fungi that are mutualists [9, 45, 46] and fungal endophytes that might protect seaweeds against pathogenic protists [44].

Adding to the complexity of the microeukaryotic biodiversity associated with brown algae, their thalli are often overgrown with a wide variety of smaller seaweeds which represents a continuum between epiphytism and endophytism. These epi-endophytes can have negative effects such as imposing physical and physiological stress on their host, or positive ecosystem effects such as increased available habitats and food for both macroscopic and microscopic life [47]. As macroalgal symbiomes comprise multi-cellular algae as epiphytes [48, 49] and potentially pathogenic endophytes [50–53] in addition to microeukaryotes, the almost complete lack of knowledge of these eukaryotes in a holobiont context/perspective is a fundamental knowledge gap which needs to be filled in order to improve our understanding of brown algal holobionts.

The overall aim of this study was to unveil the eukaryome of seven large brown algae which are key components in coastal ecosystems: *Fucus vesiculosus*, *F*. *serratus*, *Himanthalia elongata* and *Ascophyllum nodosum* (order Fucales), *Laminaria digitata* and *Saccharina latissima* (order Laminariales) and *Saccorhiza polyschides* (order Tilopteridales). We achieved this aim by using using broadly-targeted pan-eukaryotic primers and High Throughput Sequencing (HTS). This application in host environments has so far been limited due to the technical limitation of co-amplifying host DNA when using ‘universal’ eukaryotic primers [20]. However, with sufficient sequencing depth now tractable to amplify host-associated eukaryotes much more comprehensively than was previously possible, this will become increasingly established as a standard tool for investigating host-associated eukaryomes [54, 55]. We also aimed to assess whether brown algae harbour a novel or differentially structured diversity of (micro)eukaryotes, and to ascribe putative functional roles to the host-associated microeukaryotes (e.g., putative parasites).

## Results

### Composition of the brown algal eukaryome

A total of 256 eukaryotic Operational Taxonomic Units (OTUs) were defined by UPARSE and post-UPARSE trimming. Their read abundance in each library are shown in Supplementary Table S1. Each library represented five brown algal samples pooled according to species and geographic location, and is hereafter referred to as samples. The richness recovered from each sample ranged from 26 to 101 OTUs, and their taxonomic profiles are shown as proportional OTU abundance in Fig. 1, Supplementary Table S1. The sequence dataset was dominated by brown algal reads (97.3% of all sequences, Supplementary Table S2). Brown algae also displayed the highest OTU richness (18.9% of the total OTUs; Fig. 2, and accountable for 27.3-70.4% of the OTUs per sample; Supplementary Table S2). Nevertheless, there was a broad taxonomic diversity of microeukaryotes in the brown algal samples with OTUs taxonomically assigned to ten of the eukaryotic major Kingdoms or supergroups (Fig. 1, Fig. 2).

**Figure 1.**
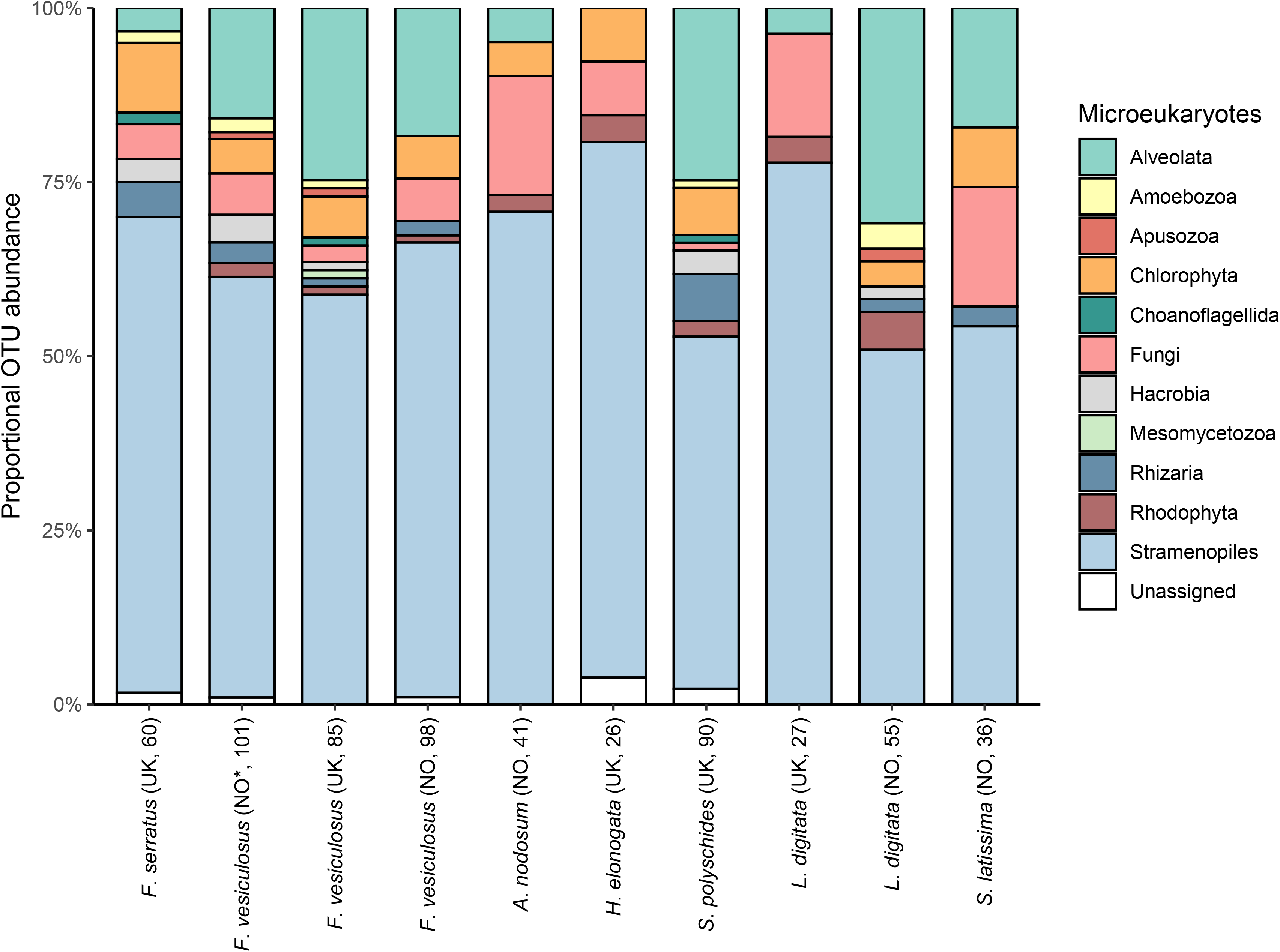
Diversity of eukaryotes in brown algal holobionts. Proportional abundance of OTUs taxonomically assigned to eukaryotic major Kingdoms or ‘supergroup’ in the brown algal samples of *Fucus serratus*, *F*. *vesiculosus*, *Ascophyllum nodosum*, *Himanthalia elongata*, *Saccorhiza polyschides*, *Laminaria digitata* and *Saccharina latissima*. The sampling location for each brown alga is shown in parenthesis; NO = Norway and UK = The United Kingdom, together with the total number of OTUs per library. The asterisk (NO*) represent *F*. *vesiculosus* sampled in Norway, May 2013. All other samples were collected in October 2015.

**Figure 2.**
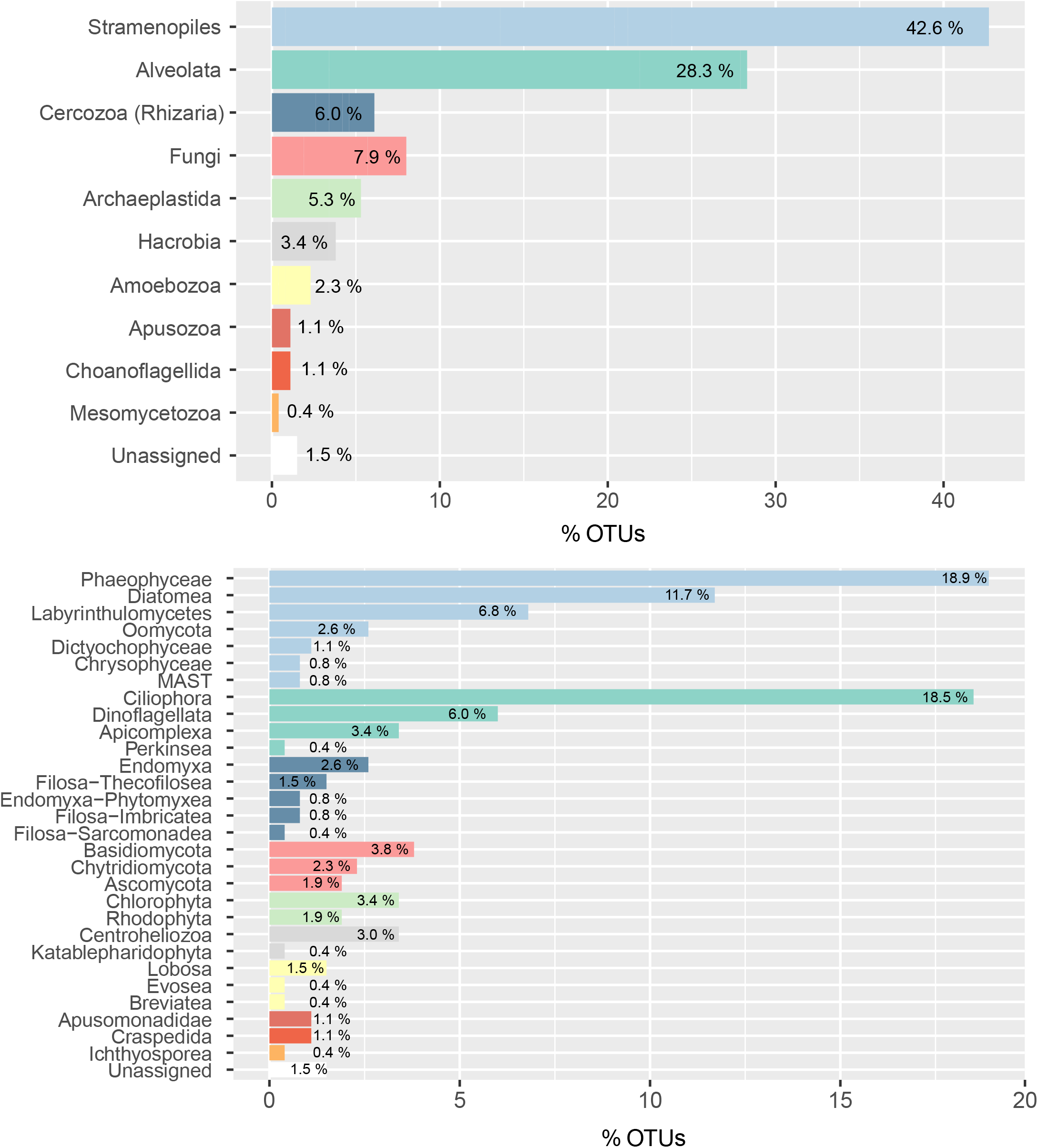
Abundance of eukaryotic taxonomic groups. Percentage representation of OTUs assigned to the different taxonomic groups at (A) Kingdom or ‘supergroup’ level and at lower taxonomic level (B) in all of the brown algal samples combined. To see the percentage of reads and percentage of OTUs for all the taxonomic groups per brown algal sample, see Supplementary table S2.

### Phylogenetic diversity of eukaryotic OTUs

Phylogenetic analyses were conducted for taxonomic groups displaying either high taxonomic diversity and/or taxonomic groups previously documented to be in a symbiotic relationship (*sensu lato*) with brown algae. The resulting phylogenies and heat maps of proportional read abundance (log_10_) of each OTU in the different samples are shown in Figs. 3-7 and Supplementary Figs. 1-4, which include OTUs and the closest sequence matches in NCBI GenBank as of 05.05.2020 (re-blasted 11.04.2021).

**Figure 3.**
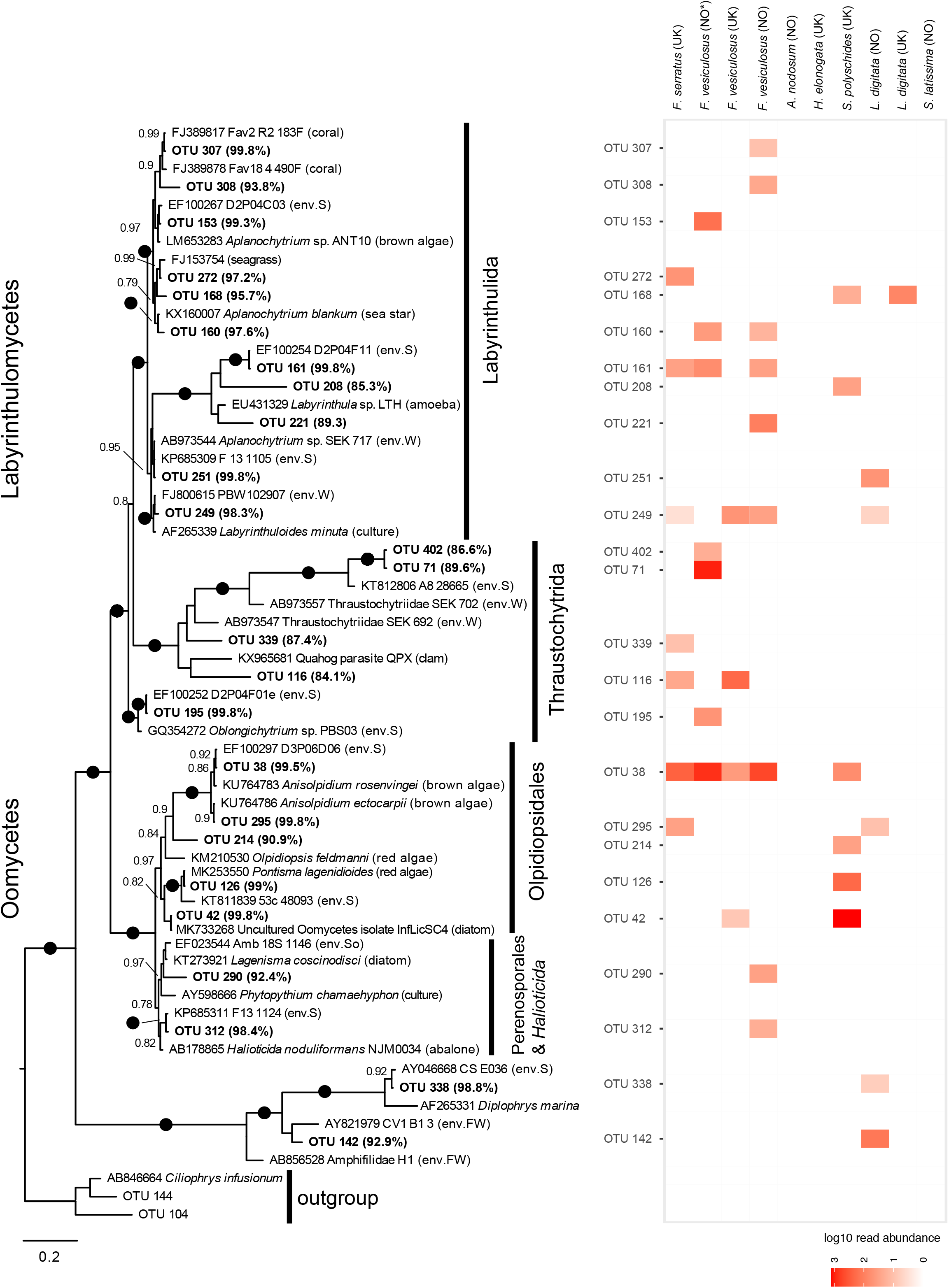
Oomycetes and labyrinthulids phylogeny with heatmap representing proportional read abundance (log_10_) of OTUs per sample. Bayesian phylogeny of oomycetes and Labyrinthulomycetes with bootstrap values from maximum likelihood analysis added. Black circles represent strong support (posterior probability >/= 0.95, and bootstrap support >/= 95%). OTUs from this study are shown in bold. The percentage identity to the most similar reference sequence is shown in parenthesis after each OTU. Host or type of environment the reference sequences were retrieved from in previous studies are listed in parenthesis after each GenBank accession number. The scale bar represents 0.2 substitutions per site. Abbreviations used for descriptions of environment: env.S = environmental sample, marine sediment; env.W = environmental sample, marine water; env.FW = environmental sample, fresh water; env.So = environmental sample, soil. The heatmap illustrates the log_10_ read abundance for each OTU in the brown algal samples of *Fucus serratus*, *F*. *vesiculosus*, *Ascophyllum nodosum*, *Himanthalia elongata*, *Saccorhiza polyschides*, *Laminaria digitata* and *Saccharina latissima*. The sampling location for each brown alga is shown in parenthesis; NO = Norway and UK = The United Kingdom. All samples were collected in October 2015, except *F*. *vesiculosus* (NO*) which was sampled in Norway, May 2013.

**Figure 4.**
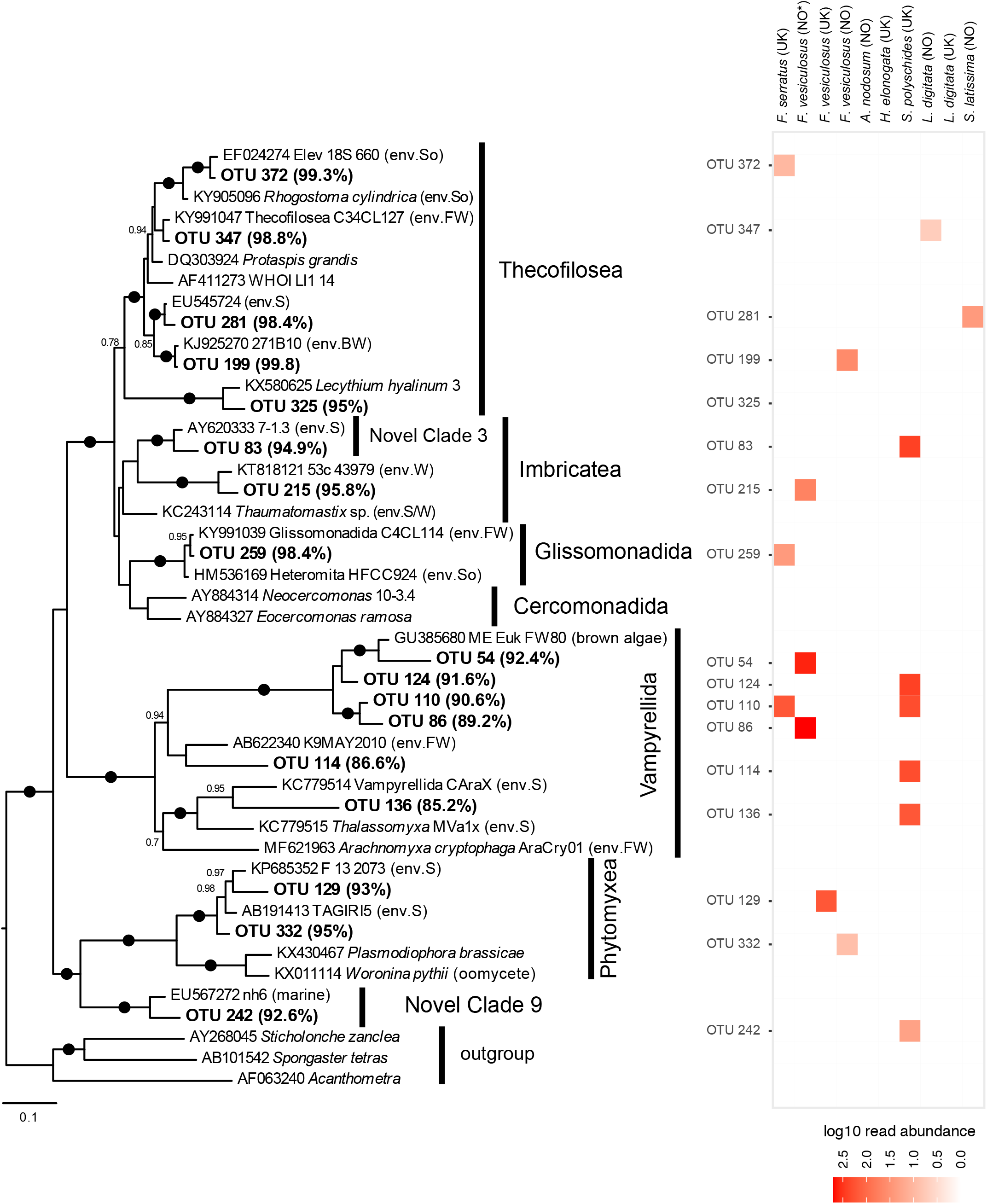
Cercozoa: phylogeny with heatmap representing proportional read abundance (log_10_) of OTUs per sample. Bayesian phylogeny of Cercozoa with bootstrap values from maximum likelihood analysis added. Black circles represent strong support (posterior probability >/= 0.95, and bootstrap support >/= 95%). OTUs from this study are shown in bold. The percentage identity to the most similar reference sequence is shown in parenthesis after each OTU. Host or type of environment the reference sequences were retrieved from in previous studies are listed in parenthesis after each GenBank accession number. The scale bar represents 0.1 substitutions per site. Abbreviations used for descriptions of environment: env.S = environmental sample, marine sediment; env.W = environmental sample, marine water; env.FW = environmental sample, fresh water; env.BW = environmental sample, brackish water; env.S/W = environmental sample, marine sediment or water; env.So = environmental sample, soil. The heatmap illustrates the log_10_ read abundance for each OTU in the brown algal samples of *Fucus serratus*, *F*. *vesiculosus*, *Ascophyllum nodosum*, *Himanthalia elongata*, *Saccorhiza polyschides*, *Laminaria digitata* and *Saccharina latissima*. The sampling location for each brown alga is shown in parenthesis; NO = Norway and UK = The United Kingdom. All samples were collected in October 2015, except *F*. *vesiculosus* (NO*) which was sampled in Norway, May 2013.

**Figure 5.**
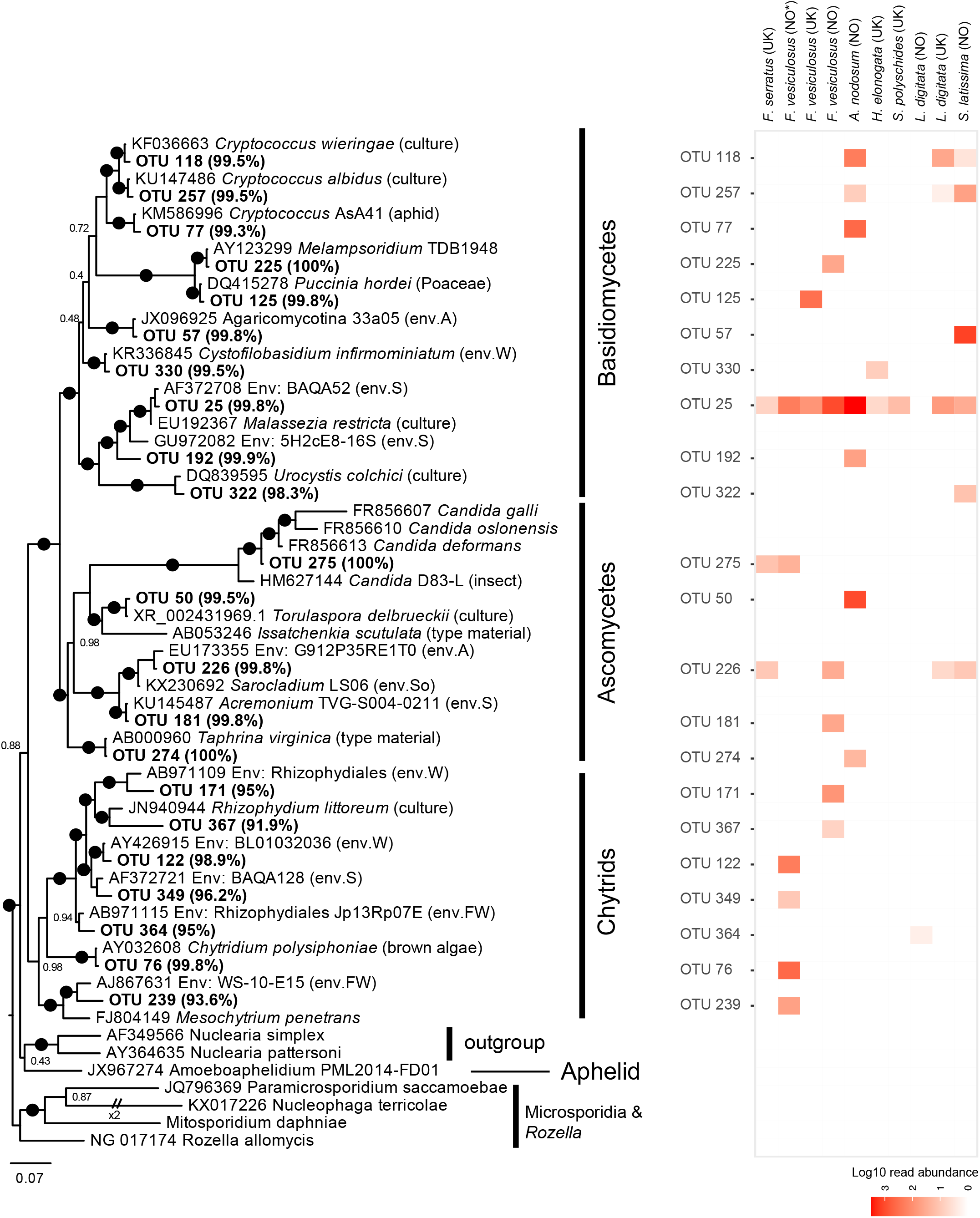
Fungi: phylogeny with heatmap representing proportional read abundance (log_10_) of OTUs per sample. Bayesian phylogeny of Fungi with bootstrap values from maximum likelihood analysis added. Black circles represent strong support (posterior probability >/= 0.95, and bootstrap support >/= 95%). OTUs from this study are shown in bold. The percentage identity to the most similar reference sequence is shown in parenthesis after each OTU. Host or type of environment the reference sequences were retrieved from in previous studies are listed in parenthesis after each GenBank accession number. The scale bar represents 0.07 substitutions per site. Abbreviations used for descriptions of environment: env.S = environmental sample, marine sediment; env.W = environmental sample, marine water; env.FW = environmental sample, fresh water; env.So = environmental sample, soil; env.A = environmental sample, air. The heatmap illustrates the log_10_ read abundance for each OTU in the brown algal samples of *Fucus serratus*, *F*. *vesiculosus*, *Ascophyllum nodosum*, *Himanthalia elongata*, *Saccorhiza polyschides*, *Laminaria digitata* and *Saccharina latissima*. The sampling location for each brown alga is shown in parenthesis; NO = Norway and UK = The United Kingdom. All samples were collected in October 2015, except *F*. *vesiculosus* (NO*) which was sampled in Norway, May 2013.

**Figure 6.**
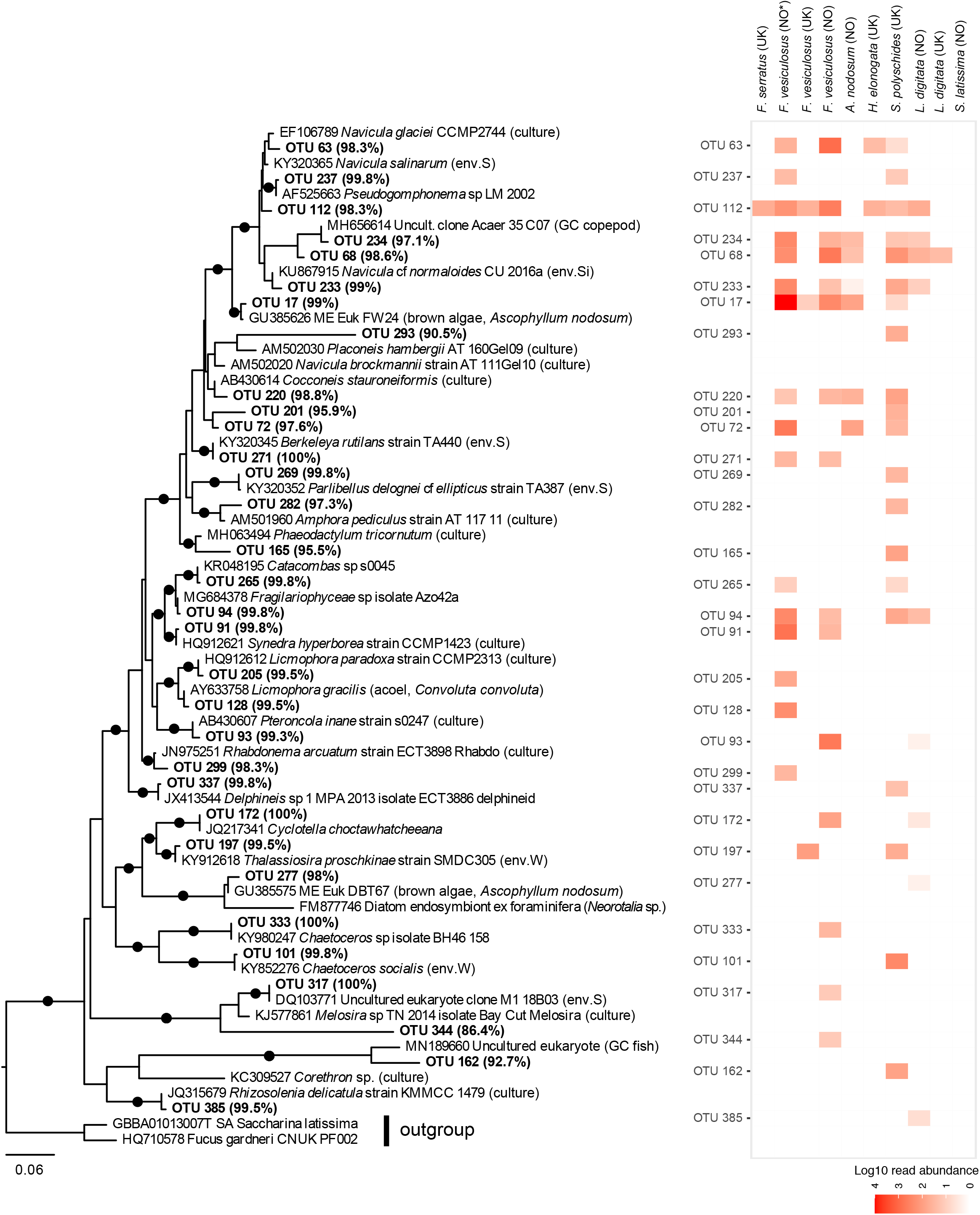
Bacillariophyceae: phylogeny with heatmap representing proportional read abundance (log_10_) of OTUs per sample. Bayesian phylogeny of diatoms with bootstrap values from maximum likelihood analysis added. Black circles represent strong support (posterior probability >/= 0.95, and bootstrap support >/= 95%). OTUs from this study are shown in bold. The percentage identity to the most similar reference sequence is shown in parenthesis after each OTU. Host or type of environment the reference sequences were retrieved from in previous studies are listed in parenthesis after each GenBank accession number. The scale bar represents 0.06 substitutions per site. Abbreviations used for descriptions of environment: env.S = environmental sample, marine sediment; env.W = environmental sample, marine water; env.Si = environmental sample, sea ice; GC = gut content. The heatmap illustrates the log_10_ read abundance for each OTU in the brown algal samples of *Fucus serratus*, *F*. *vesiculosus*, *Ascophyllum nodosum*, *Himanthalia elongata*, *Saccorhiza polyschides*, *Laminaria digitata* and *Saccharina latissima*. The sampling location for each brown alga is shown in parenthesis; NO = Norway and UK = The United Kingdom. All samples were collected in October 2015, except *F*. *vesiculosus* (NO*) which was sampled in Norway, May 2013.

**Figure 7.**
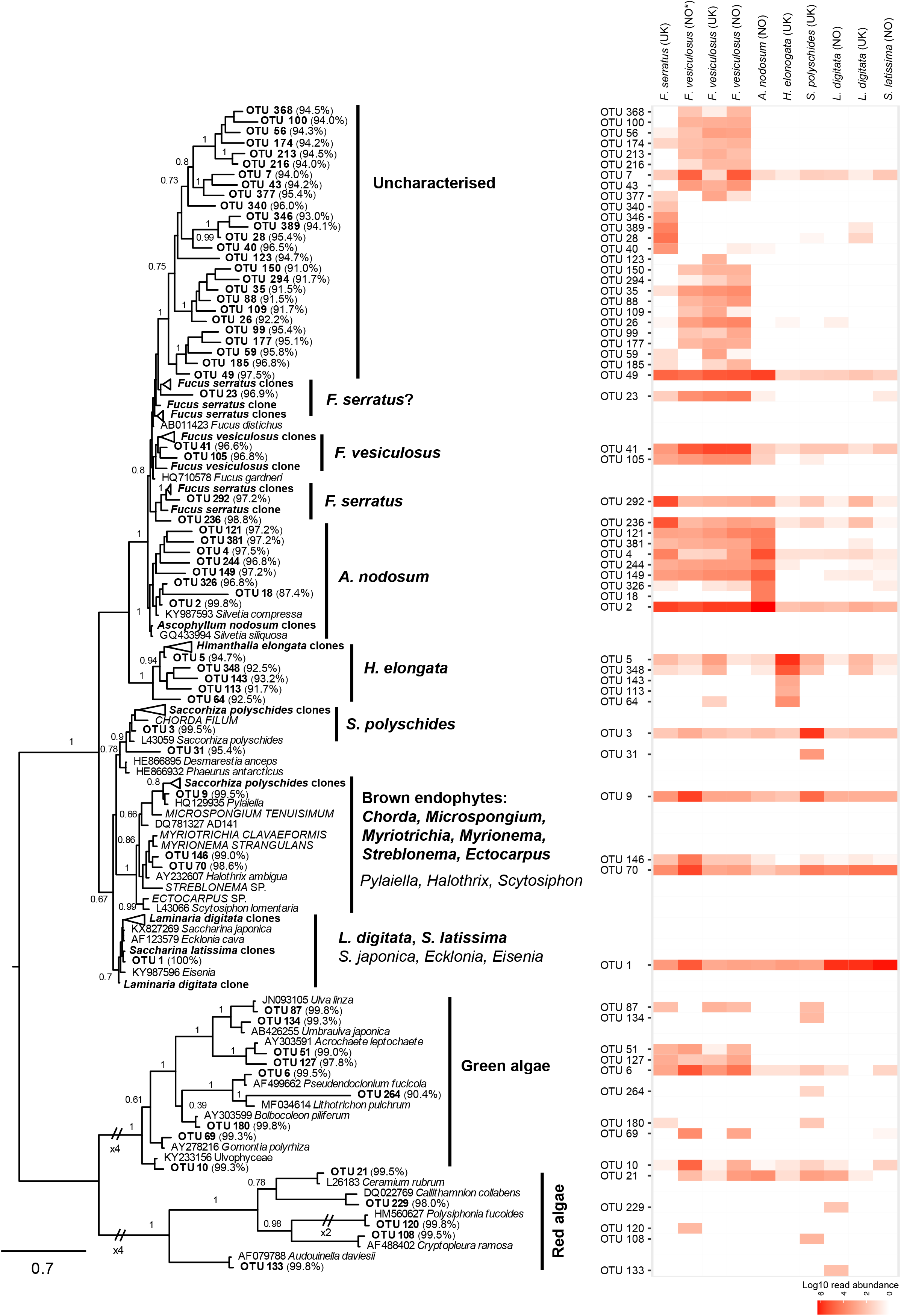
Brown-Red-Green: phylogeny with heatmap representing proportional read abundance (log_10_) of OTUs per sample. Bayesian phylogeny of Phaeophyceae, Rhodophyta and Chlorophyta with bootstrap values from maximum likelihood analysis added. Black circles represent strong support (posterior probability >/= 0.95, and bootstrap support >/= 95%). OTUs from this study are shown in bold. The percentage identity to the most similar reference sequence is shown in parenthesis after each OTU. Host or type of environment the reference sequences were retrieved from in previous studies are listed in parenthesis after each GenBank accession number. The longer 18S sequences generated by cloning and sanger sequencing in this study are shown in bold and includes which brown algal host species they were amplified from. The brown algal endophytes isolated and sequenced by Attwood (2005, [141]) are all in the ‘brown endophytes’ clade and are shown in capital letters. The scale bar represents 0.7 substitutions per site. The heatmap illustrates the log_10_ read abundance for each OTU in the brown algal samples of *Fucus serratus*, *F*. *vesiculosus*, *Ascophyllum nodosum*, *Himanthalia elongata*, *Saccorhiza polyschides*, *Laminaria digitata* and *Saccharina latissima*. The sampling location for each brown alga is shown in parenthesis; NO = Norway and UK = The United Kingdom. All samples were collected in October 2015, except *F*. *vesiculosus* (NO*) which was sampled in Norway, May 2013.

For some taxonomic groups, the heatmaps indicated that the majority of OTUs were only detected associated with one brown algal sample, such as Cercozoa (Fig. 4), Fungi (Fig. 5), Ciliophora (Supplementary Fig. 1), Apicomplexa and Perkinsea (Supplementary Fig. 2), Dinoflagellata (Supplementary Fig. 4) and Amoebozoa and Centroheliozoa (Supplementary Fig. 3). However, most of these taxonomic groups also comprised OTUs that were found in multiple samples, across host orders and geographic locations (e.g., the fungal ascomycete OTU 226; Fig. 5, the ciliate OTU 89; Supplementary Fig. 1, and the centroheliozoan OTU 13; Supplementary Fig. 3 were found in samples belonging to Fucales, Tilopteridales and Laminariales from both Norway and the UK). For other taxonomic groups such as Bacillariophyta (Fig. 6) and Labyrinthulomycetes & oomycetes (Fig. 3), the heatmaps show that a large proportion of the OTUs were detected in multiple samples.

Several OTUs clustered with reference sequences that have been found associated with various hosts in previous studies, such as the labyrinthulid and oomycete OTUs clustering with *Aplanochytrium* (Labyrinthulida) and Olpidiopsidales (Fig. 3), and the cercozoan vampyrellid OTUs clustering with a reference sequence previously detected in samples of brown alga (Fig. 4).

### Novel diversity in the brown algal eukaryome

We found that ∼30% of the OTUs had low percentage identity (<95 %) with any known close relatives in reference sequence databases and represent potential novel taxonomic diversity (Supplementary Table 1). The majority of OTUs taxonomically and phylogenetically placed within Fungi, Bacillariophyta and ciliates displayed high percentage identity (>95 %) to known reference sequences (Fig. 5; Fig. 6; and Supplementary Fig. 1, respectively). Conversely, within Labyrinthulomycetes and oomycetes (Fig. 3), Cercozoa (Fig. 4) and Amoebozoa and Centroheliozoa (Supplementary Fig. 3), about 50% of the OTUs had low similarity to known reference sequences (<95 %) which indicated that these OTUs represented potential novel diversity. Some of this potential novel diversity was represented by specific clades in the phylogenetic trees, such as a well-supported labyrinthulid clade within Thraustochytrida that encompassed four OTUs with very low percentage identity (ranging between 84.1% and 89.6%). Similarly, within Cercozoa, eight of the nine Vampyrellida, Phytomyxea, and Novel Clade 9 OTUs had low similarity to reference sequences (from 85.2% to 93%; Fig 4).

### Green (Chlorophyta), red (Rhodophyta) and Brown (Phaeophyceae) algae

In addition to protist taxa, we also detected OTUs from green, red, and (non-host-derived) brown algae (Fig. 7). The majority of the nine green algal OTUs (Fig. 7) were very similar (>99% identity) to characterized and sequenced Ulvales taxa such as *Ulvella* (formerly *Acrochaete*) *leptochaete* and *Umbraulva japonica*. Only a single green alga (OTU 264) displayed low similarity (90.4%) to reference sequences, and clustered with *Lithotrichon pulchrum*, Ulvales (Fig. 7). Similarly, the five red algal OTUs (Fig. 7) displayed >98% similarity to taxa including *Ceramium* sp. and *Cryptopleura ramosa*.

There were in total 50 brown algal OTUs. Significant levels of microdiversity in the V4 OTUs were seen in the phaeophyte clade. OTUs corresponding to the host species were inferred on the basis of their phylogenetic proximity to, and high proportions of reads associating with, reference sequences from that host, either from GenBank or sequences generated in this study. Host OTUs are indicated by labeled brackets on Fig. 7. Many of the OTUs derived from the host species were present in samples from multiple host species, albeit generally represented with lower read abundance in the ‘non-host’ samples.

However, other OTUs did not cluster with host-derived sequences, or from the longer 18S sequences generated by cloning amplicons from the host. The 26 OTUs in the “Uncharacterised clade” on Fig. 7 were clearly distinct from any characterized or environmental sequences in reference sequence databases. This clade encompassed OTUs that were mainly detected in the *Fucus* sp. samples, but six of them were also detected in lower abundance in other brown algal samples (Fig. 7). Additionally, seven OTUs in the *A. nodosum* clade may be host sequences but are obviously different in sequence from the cloned sequences generated from that host, and OTUs 2 and 326, presumably deriving from that host (Fig. 7).

Some OTUs grouped strongly with known epiphytic lineages, such as the three OTUs clustering with the genera *Pylaiella* and *Halothrix*. These OTUs were found in all, or in the majority, of the different brown algal hosts (Fig. 7). In the same clades were the brown algal endophytes isolated and sequenced by Attwood (2005, [141]), published here for the first time, and represented on Fig. 7, all in the ‘brown endophytes’ clade. These endophytes were identified as *Myrionema strangulans*, isolated from *Fucus serratus*, *Mastocarpus stellatus*, *Chorda filum*, *Osmundea pinnatifida*, and *Chondrus crispus*; *Microspongium tenuisimum*, isolated from *Osmundea osmunda* and *C. crispus*; *Streblonema* sp. isolated from *M. stellatus*; *Mytriotrichia clavaeformis* and *Ectocarpus* sp. from *C. crispus*; and *Chorda filum* from a larger *C. filum* individual. Light microscopy images of the isolated endophytes are shown in Fig. 8.

**Figure 8.**
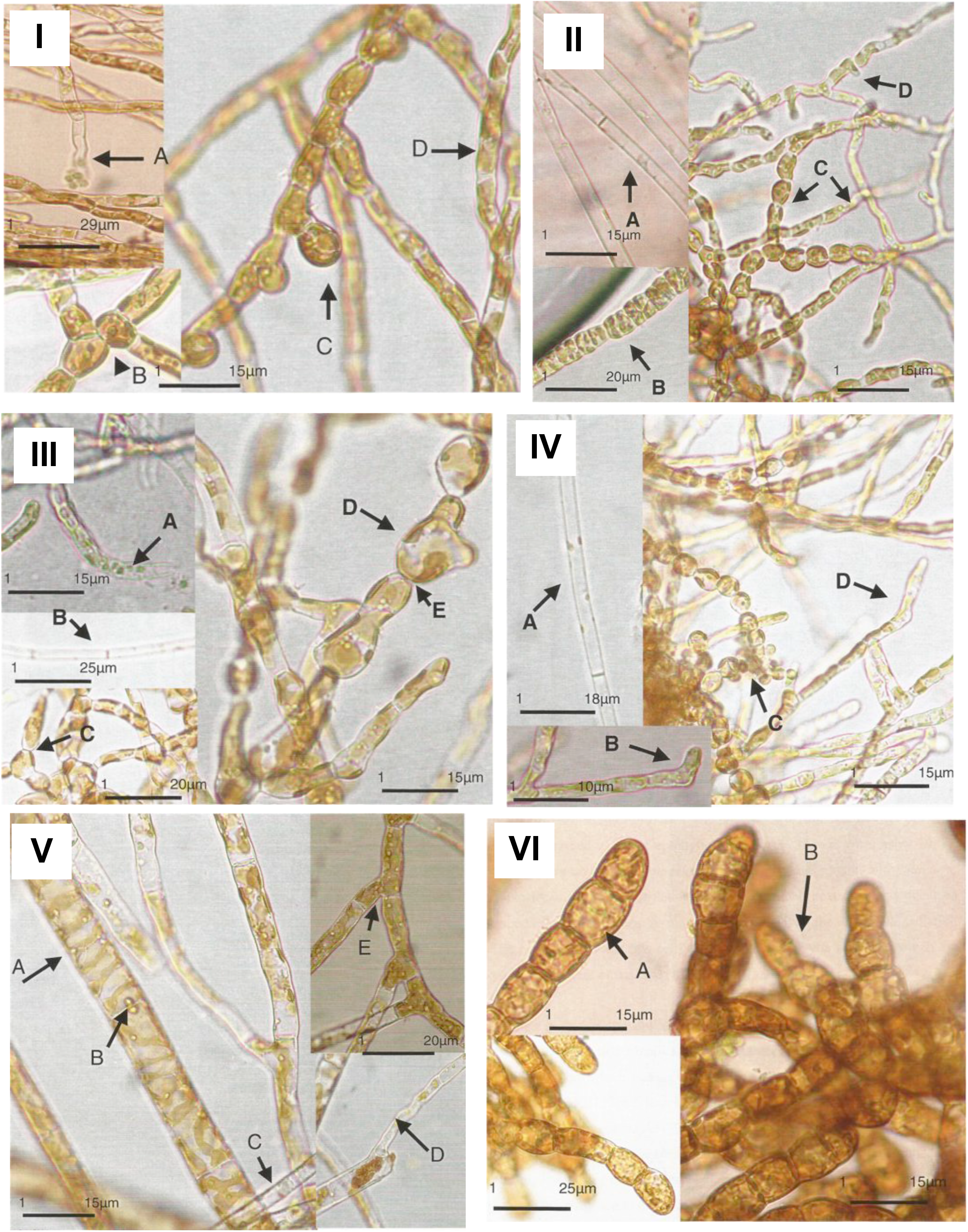
Brown algal endophytes. Light microscopy images of phaeophyte endophytes isolated from *Fucus serratus*, *Mastocarpus stellatus*, *Chorda filum*, *Osmundea pinnatifida*, and *Chondrus crispus*. (I) *Streblonema* sp.: A – terminal sporangium, showing apical spore release; B – branching thallus from spherical cells; C – growth pattern of new filament; D – discoid parietal plastid. (II) *Myrionema strangulans*: A – Phaeophyceae hair; B – plurilocular sporangia; C-two different cell shapes; D – unilocular lateral sporangium. (III) *Microspongium tenuisimum*: A - plurilocular sporangia releasing zoospores; B - Phaeophyceae hair; C – bifurcating branch; D ovate cell, showing plate-like plastid with prominent pyrenoids; E – waisted cross cell wall. (IV) *Mytriotrichia clavaeformis*: A - Phaeophyceae hair; B – clubbed terminal cell; C - unilocular clustering sporangia; D – empty clubbed terminal sporangium. (V) *Ectocarpus* sp.: A – ribbon-shaped chloroplast; B – pyrenoid; C – pseudohairs; D – pseudohairs shoing growth from filament; E – branching growth form. (VI) *Chorda filum*: A – densely pigmented cells; B prostrate clumps of branched filaments.

Three of our OTU sequences occurred in the same clade as these endophytes: OTU 9 and some 18S clone sequences from *S. polyschides* clustering with *Pylaiella*, and OTUs 70 and 146, clustering most closely with *Myriotrichia* and *Myrionema* (Fig. 7). All of these were strongly associated with *F. vesiculosus* and *S. polyschides* individuals, but they were present in almost all host samples.

### Analyses of community composition

The clustering of samples was consistent irrespective of whether brown algal OTUs were included in the analyses (Fig 9: A-B). The three *F*. *vesiculosus* samples clustered with *F*. *serratus* and *S*. *polyschides* in both plots (Fig 9: A-B), which indicated that these have a similar OTU composition. The *A*. *nodosum* sample showed less similarity with the *Fucus* and *S*. *polyschides* samples when the brown algal OTUs were excluded from the analysis (Fig. 9, A-B). *Himanthalia elongata* did not cluster with any other brown algal hosts and appeared highly dissimilar from the others both including and excluding brown algal OTUs (Fig. 9; A-B). Analyses of similarity (ANOSIM) were highly significant both with and without brown algal OTUs included (p-value 7e-04 and 0.03 respectively), which supported the dissimilarity observed between the brown algal host genera in the NMDS plots (Fig. 9: A-B).

**Figure 9.**
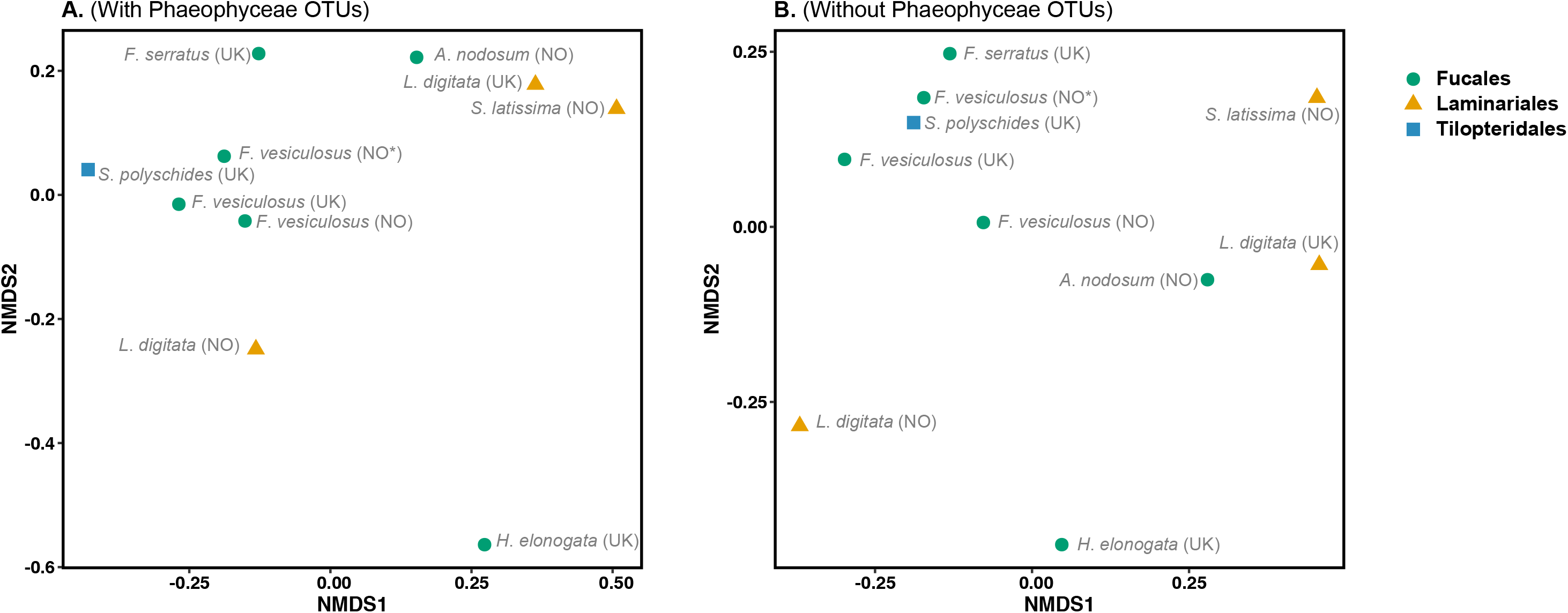
Community composition. Non metric multidimensional scaling (NMDS) analysis of the composition of eukaryotic OTUs in the brown algal samples based on Bray-Curtis pairwise distances. (A) including brown algal/Phaeophyceae OTUs and (B) excluding brown algal/Phaeophyceae OTUs.

Dissimilarity between host genera (SIMPER values) ranged from 31.4% (*Laminaria* spp. vs *Saccharina* spp.) to 99.9% (*Ascophyllum* spp. vs *Himanthalia* spp.) in the dataset with brown algal OTUs included (Table 1, Supplementary Table S3). Certain host derived OTUs (OTU 1, 2, 3 and 5) were the main contributors to the observed dissimilarity. These results also supported some of the OTUs inferred to derive from the host species based on their phylogenetic placements and read abundances (Table 1, Fig. 7). When brown algal OTUs were excluded the host genera were all highly dissimilar, with SIMPER values ranging from 83.3% (*Ascophyllum* spp. vs *Saccorhiza* spp.) to 99.4% (*Ascophyllum* spp. vs *Himanthalia* spp.). The main contributors to the observed dissimilarity between the brown algal host genera were green algal, red algal, fungal and oomycete OTUs in addition to an unassigned OTU (Table 2). There were also several OTUs assigned to a wide range of taxonomic groups that had some contribution to the dissimilarity (Supplementary Table S3).

**Table 1.** Summary of pairwise SIMPER analyses between brown algal host samples grouped at genus level. Showing the OTUs that contributed to more than 10% of the observed dissimilarity between host genera according to SIMPER, when Phaeophyceae OTUs were included in the analyses.

|  |  | OTUs contributing >10 % to the dissimilarity between brown algal hosts<br>(Including Phaeophyceae OTUs) |  |  |  |  |  |
| --- | --- | --- | --- | --- | --- | --- | --- |
|  | Phylum | Stramenopiles |  |  |  |  |  |
|  | Class | Phaeophyceae |  |  |  |  |  |
|  | PR2 taxonomy | *Silvetia siliquosa* | *Fucus gardneri* | *Fucus gardneri* | *Silvetia siliquosa* | *Saccorhiza polyschides* | *Agarum clathratum* |
|  | Inferred host taxonomy | *Ascophyllum nodosum* | *Fucus spp.* | *Fucus vesiculosus* | *Himanthalia elongata* | *Saccorhiza polyschides* | *Laminariales*<br>(*L. digitata* + *S. latissima*) |
| SIMPER comparison | Dissimilarity (%) | OTU 2 | OTU 49 | OTU 41 | OTU 5 | OTU 3 | OTU 1 |
| Fucus – Ascophyllum | 74.4 | 41.3 | 8.4 | 4.8 |  |  |  |
| Fucus – Himanthalia | 99.8 | 20.0 | 5.1 | 9.6 | 36.6 |  |  |
| Fucus – Saccorhiza | 97.9 | 21.1 | 5.4 | 10.1 |  | 35.8 |  |
| Fucus – Laminaria | 96.7 | 21.0 | 5.4 | 10.1 |  |  | 38.3 |
| Fucus – Saccharina | 97.3 | 15.2 | 3.9 | 7.4 |  |  | 54.3 |
| Ascophyllum – Himanthalia | 99.9 | 54.9 | 11.7 |  | 19.7 |  |  |
| Ascophyllum – Saccorhiza | 99.8 | 56.6 | 12.1 |  |  | 18.7 |  |
| Ascophyllum – Laminaria | 99.9 | 56.4 | 12.1 |  |  |  | 20.8 |
| Ascophyllum – Saccharina | 99.9 | 47.3 | 10.1 |  |  |  | 33.3 |
| Himanthalia – Saccorhiza | 99.7 |  |  |  | 45.0 | 41.6 |  |
| Himanthalia – Laminaria | 99.7 |  |  |  | 44.7 |  | 45.8 |
| Himanthalia – Saccharina | 99.8 |  |  |  | 31.3 |  | 61.6 |
| Saccorhiza – Laminaria | 98.3 |  |  |  |  | 44.2 | 49.1 |
| Saccorhiza – Saccharina | 98.5 |  |  |  |  | 30.4 | 64.6 |
| Laminaria – Saccharina | 31.4 |  |  |  |  |  |  |

**Table 2.**
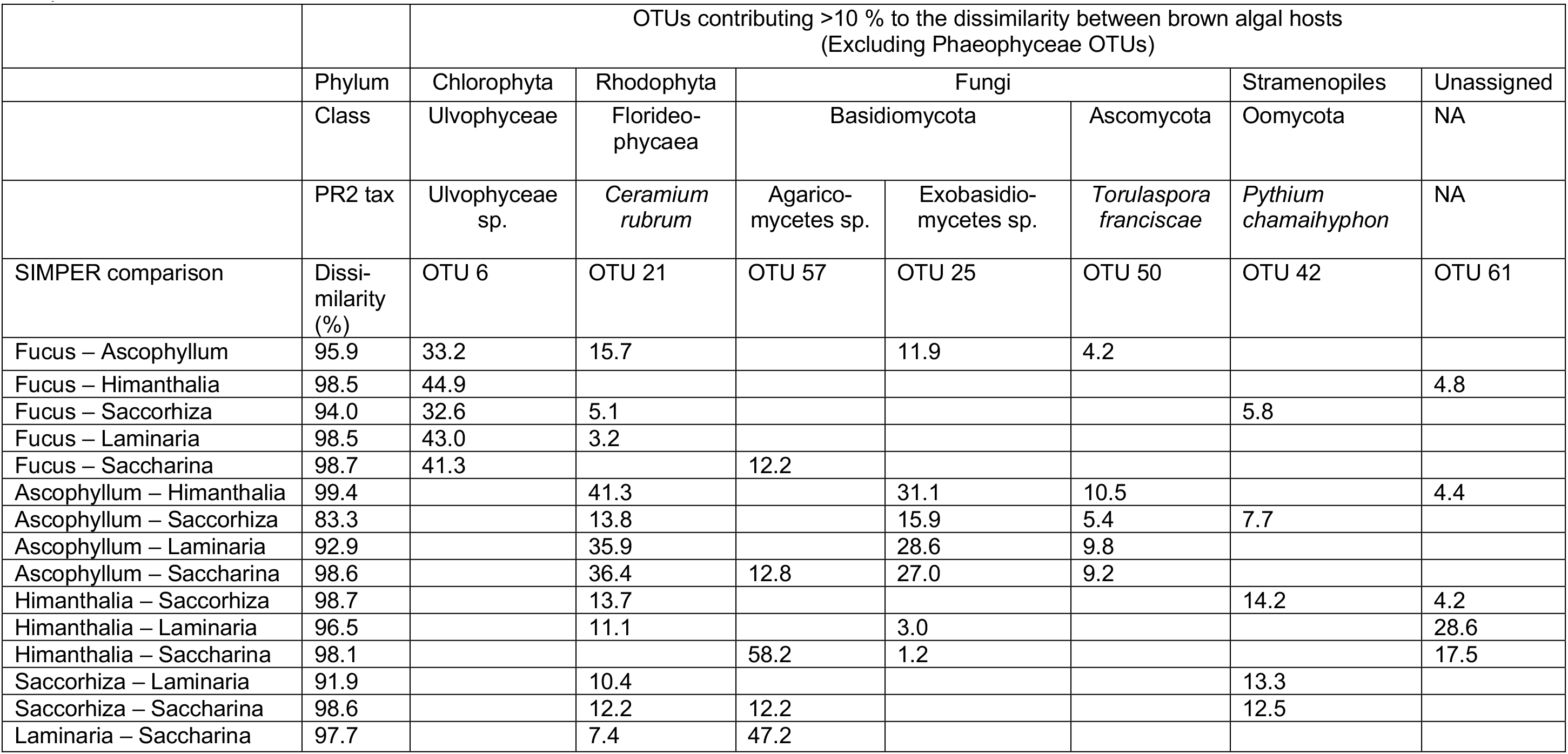
Summary of pairwise SIMPER analyses between brown algal host samples grouped at genus level. Showing the OTUs that contributed to more than 10% of the observed dissimilarity between host genera according to SIMPER, when Phaeophyceae OTUs were excluded from the analyses.

IndVal analyses of the dataset including brown algal OTUs, indicated that six OTUs were preferentially associated with *Ascophyllum* (Table 3), of which five were taxonomically assigned to Phaeophyceae. Four of these OTUs clustered within the clade labelled “*A*. *nodosum*” in the phylogenetic tree (Fig. 7, Table 3), while the fifth OTU was placed in the “Uncharacterised” clade, represented with reads in all the brown algal hosts (Fig. 7). The final OTU (25) was a fungal OTU found in all samples except *L*. *digitata* (UK), but with higher read abundance in *Ascophyllum* than in the other host genera (Fig. 5). OTU 25 was also the only one found to be significant by the IndVal analysis of the dataset excluding brown algal OTUs (p-value 0.0446; Table 2).

**Table 3.** Indicator species analysis (IndVal) showing OTUs most preferentially associated with the brown algal hosts.

| OTU# | Associated with | p-value | Taxonomic assignment |
| --- | --- | --- | --- |
| OTU 149 | *Ascophyllum* | 0.0185 * | Stramenopiles; Ochrophyta; Phaeophyceae; *Silvetia*; *Silvetia siliquosa* |
| OTU 381 | *Ascophyllum* | 0.0175 * | Stramenopiles; Ochrophyta; Phaeophyceae; *Silvetia*; *Silvetia siliquosa* |
| OTU 244 | *Ascophyllum* | 0.0005 *** | Stramenopiles; Ochrophyta; Phaeophyceae; *Silvetia*; *Silvetia siliquosa* |
| OTU 2 | *Ascophyllum* | 0.0020 ** | Stramenopiles; Ochrophyta; Phaeophyceae; *Silvetia*; *Silvetia siliquosa* |
| OTU 49 | *Ascophyllum* | 0.0127 * | Stramenopiles; Ochrophyta; Phaeophyceae; *Fucus*; *Fucus gardneri* |
| OTU 25 | *Ascophyllum* | 0.0446 * | Fungi; Basidiomycota; Ustilaginomycotina; Exobasidiomycetes; Exobasidiomycetes_X sp. |
| OTU 1 | *Laminaria* + *Saccharina* | 0.0033 ** | Stramenopiles; Ochrophyta; Phaeophyceae; *Agarum*; *Agarum clathratum* |

The Associated with column show which library/host species the OTUs were preferentially associated with. Abbreviations used; *Ascophyllum* = *Ascophyllum nodosum*, *Laminaria* = *Laminaria digitata*, *Saccharina* = *Saccharina latissima*. The asterisk associated with IndVal p-values indicate significance at $\leq 0.001$ (\*\*\*), $\leq 0.01$ (\*\*), $\leq 0.05$ (\*).

### Shared OTUs

Although our analyses suggested a large degree of dissimilarity between the different brown algal hosts (Table 1, 2), 70 OTUs were found in multiple samples across the host orders Fucales, Laminariales and Tilopteridales (Fig. 10, Table 4). Of these, 25 OTUs were detected in samples belonging to all of the three host orders, of which 15 belonged to Phaeophyceae, including the three brown algal endophyte OTUs (Fig. 7), three OTUs from the “Uncharacterised” clade (Fig. 7) and several host OTUs that were found in high abundances in the respective host samples, but also present in lower abundance in several of the other samples. The shared OTUs also included five diatoms which clustered with known epiphytes and endophytes of macroalgae in the phylogenetic tree (Fig. 6, Table 4).

**Figure 10.**
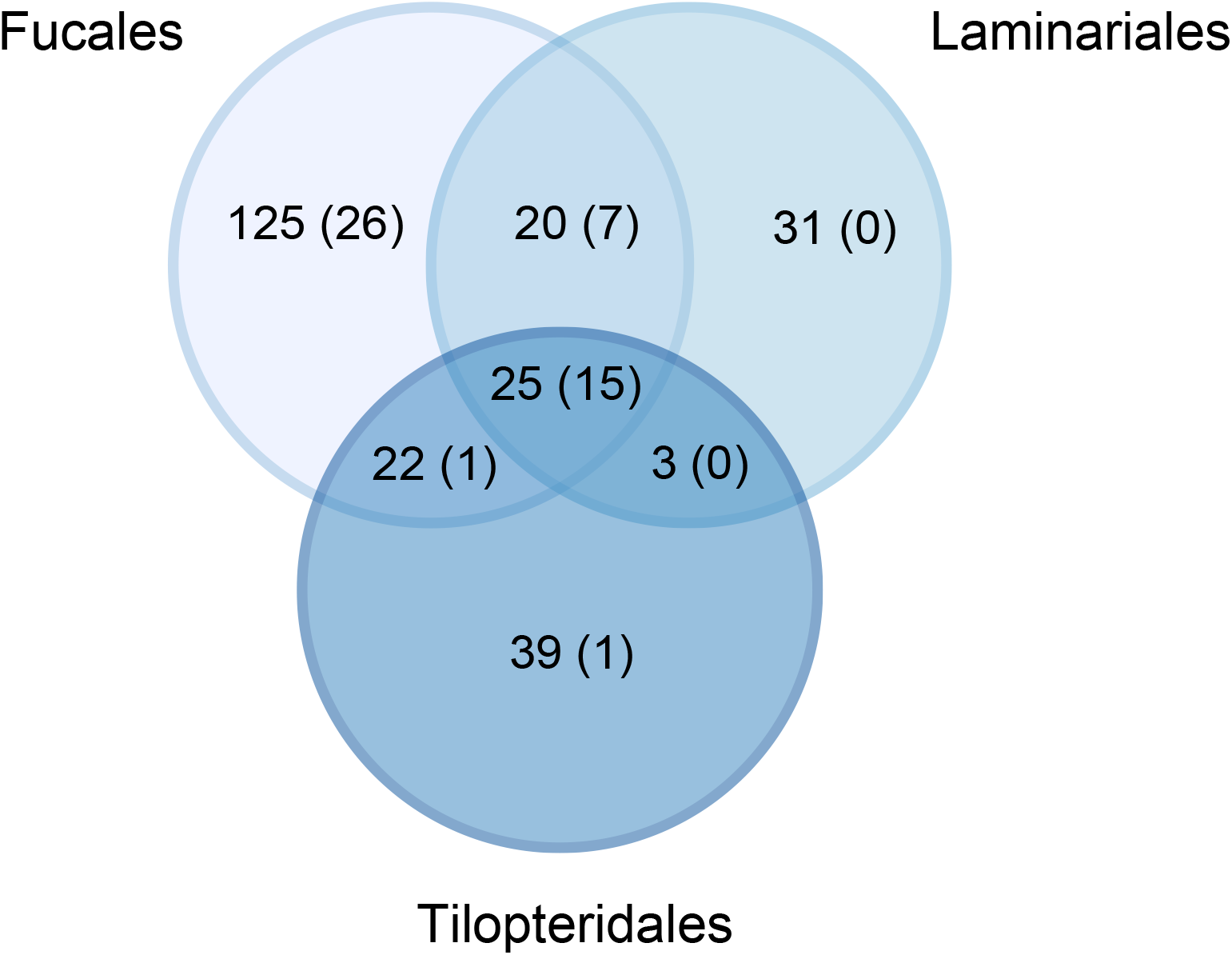
Shared and unique OTUs in the brown algal orders. Venn diagrams displaying the number of shared and unique OTUs between the brown algal host orders Fucales, Laminariales and Tilopteridales. Numbers in parentheses show the number of phaeophyte OTUs in the shared/unique data. To see the taxonomic assignment for the shared OTUs see Table 4, and for the unique OTUs see Supplementary Table S4.

**Table 4.**
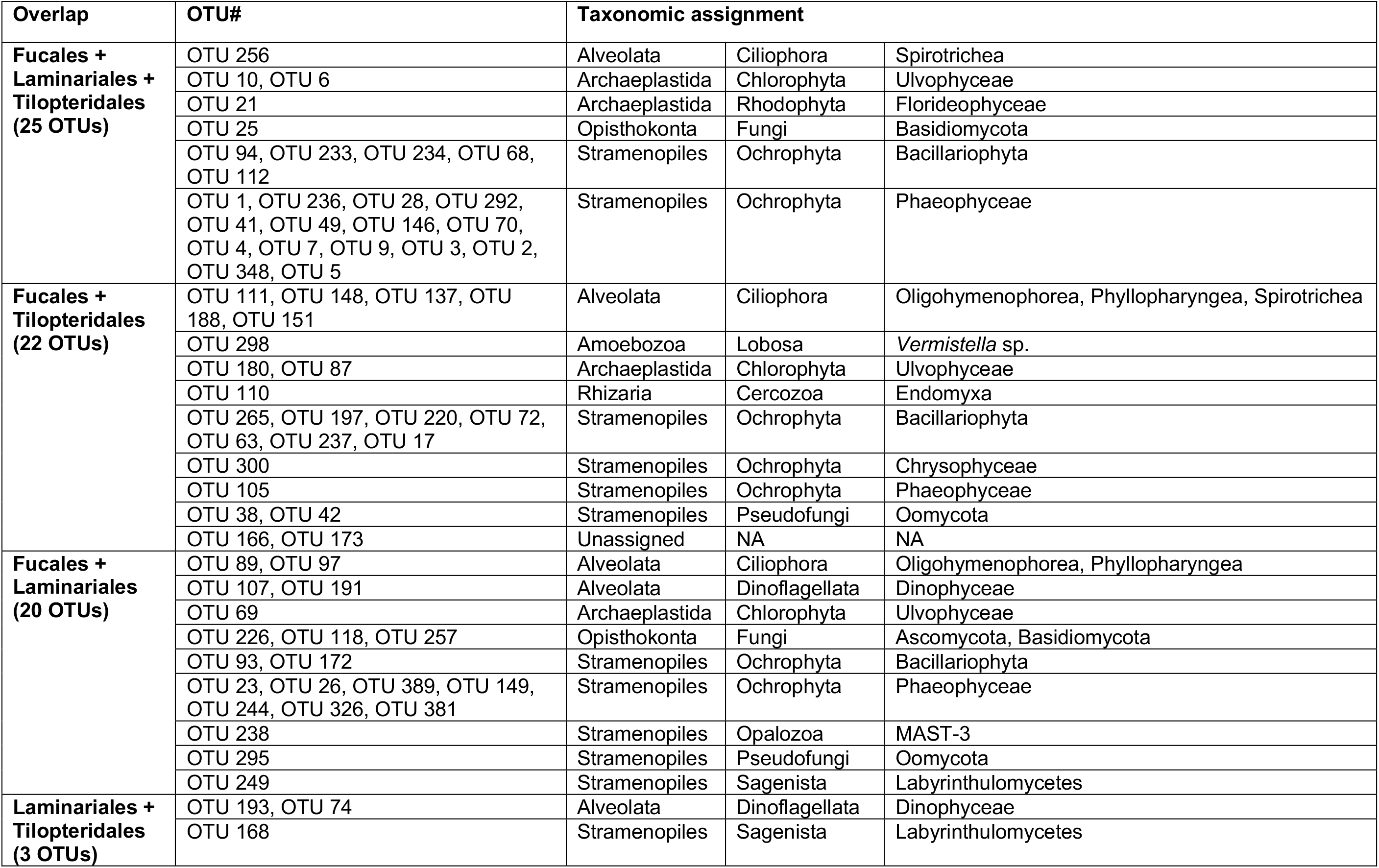
Taxonomy of the shared OTUs between the three brown algal orders: Fucales, Laminariales and Tilopteridales, as shown in Venn plot (Fig. 10).

Twenty OTUs were found in samples belonging to Fucales and Laminariales (Table 4), including Fungi, oomycetes and Labyrithulomycetes (Table 4, Fig. 5, 3). The 22 OTUs that overlapped between Fucales and Tilopteridales comprised several diatom and green algal OTUs (Table 4, Fig 6, 7), and two oomycete OTUs that clustered within Olpidiopsidales (Table 4, Fig. 3). The OTUs that were unique to Fucales, Tilopteridales and Laminariales are shown in Supplementary Table S4.

## Discussion

### Broad diversity of microeukaryotes associated with brown algae

To our knowledge, this is the first study that has investigated the diversity of eukaryotes associated with brown algae using high throughput sequencing. Our findings show that, although the sequence data was dominated by brown algal reads, we were able to identify a broad diversity of microeukaryotes representing most of the main branches in the eukaryotic tree of life (Fig 1, 2).

Some of the brown algae we investigated displayed a higher diversity of microeukaryotes compared to others. Particularly, we detected a higher diversity associated with *Fucus vesiculosus*, *F*. *serratus* (Fucales) and *Saccorhiza polyschides* (Tilopteridales) than *Laminaria digitata* and *Saccharina latissima* (Laminariales). Density and composition of microorganisms associated with brown algae have been shown to vary at several scales: e.g., among host species, between individuals of the same host species, different thallus areas of individual hosts, the kind of surface tissue available for microepiphyte colonization, different habitats and location in the intertidal zone, seasons, different stages of the life cycle, and different longevity of the host [e.g., 19, 25, 32, 56, 57-59]. In addition, the interaction between the settlers affects the formation of specific communities [47, 56], including chemical interactions and signalling between the microbial symbionts [44, 60, 61]. Seaweeds themselves also, to various degrees have physical and chemical antifouling defence mechanisms which prevent settlement of microorganisms and epiphytes [31, 62, 63]. For this study, we may also have randomly sampled different microhabitats of the brown algal thalli, resulting in a highly variable diversity of microeukaryotes.

The community composition of microeukaryotes associated with *Himanthalia elongata* stood out as very dissimilar to the other phaeophytes (Fig. 9 B). This seaweed differs from the other brown algae in the type of life cycle and the tissue type sampled for this study. *Himanthalia elongata* is biannual, but the elongated thongs which accounts for ∼98% of the thallus is reproductive tissue that degrades at the end of the season (reproductive season from late July to ∼January), leaving only the vegetative basal disc [64]. The tissue type sampled from *H*. *elongata* were therefore different and most likely younger than the tissues sampled from the other brown algae, which could explain the distinctive and low diversity of microeukaryotes detected in this host. A recent study also reported low diversity of bacteria associated with *H*. *elongata* [65].

### Eukaryotic epiphytes and endophytes in the seaweed holobiont

Seaweed surfaces provide nutrient-rich and important habitats for epiphytic and endophytic organisms, from unicellular microbes such as bacteria and protists to multicellular, usually filamentous seaweeds [e.g., 10, 12, 47, 56, 66]. These associations can be either facultative (i.e. the epi-endophytes can grow on a variety of biotic and/or abiotic substrata) or obligate (dependent on one or more specific hosts). Like many biological categorizations, the boundaries between epiphytes and endophytes in real-life can be blurred; the nature of the association can be affected by both biotic and abiotic factors, and can range from mutualism to parasitism [47, 67, 68].

### Microeukaryotic epi-endophytes

Diatoms are regularly found in the biofilm on seaweed surfaces, where they live as epiphytes, and often occur in large numbers [32, 69–71]. Although many diatoms associated with brown algae are epiphytes, there are also studies which have demonstrated that some diatom species, such as the ones belonging to the genera *Navicula* and *Cocconeis* can live as endophytes, intracellularly in tissues of both brown algae and red algae [30, 72–74]. Mayombo *et al*. [75, 76] recently suggested that some diatom taxa might have adapted to an endophytic life to avoid the antifouling mechanisms of their hosts. Several of the OTUs from our study which were taxonomically and phylogenetically assigned to different species of Naviculales, Cocconeidales and Fragilariales (Fig. 6), were present in multiple samples/hosts independent of geographic location. Our molecular data in combination with previous observational studies [30, 32, 72–76] supports the hypothesis that some diatom species such as *Navicula* sp. and *Cocconeis* sp., might have a more intimate, potentially endophytic, association with seaweeds.

Ciliates represent another ubiquitous group of microeukaryotes living as epiphytes in the biofilm on seaweeds [8, 77], and was also the dominating group of alveolates in our study (Fig. 2). Many ciliates are predators/grazers feeding on bacteria and microeukaryotes [8, 78], whereas other ciliates can be involved in symbiotic interactions [78, 79]. Most of the ciliates detected in our study were only found in single samples (Supplementary Fig. 1), which might indicate that these ciliates were “random visitors” without any specific interaction with brown algae. A few of the OTUs were, however found in several samples from both Norway and the UK, and were highly similar to reference sequences from different marine environmental samples. This suggests that these ciliates are generalist biofilm formers on different organisms and substrates. However, one of these OTUs (OTU 256, Spirotrichea, Supplementary Fig. 1) was almost identical to a reference sequence found in samples of *Ascophyllum nodosum* used as live bait wrapping [80], which could indicate that some of these ciliates have a closer association with brown algae. As emphasised by del Campo et al. [20], “it is likely that many microeukaryotes routinely detected or observed/cultured from environmental samples may also be host associated, even if transiently”.

### Host-associated phaeophytes, chlorophytes and rhodophytes

In addition to the brown algal OTUs clearly deriving from the host species, we detected a diversity of other sequence types from putative brown algal epi-endophytes (Fig. 7). The diversity of ‘OTUs’ that apparently derived from the host organisms was much higher than expected. This may partly be caused by artefactual sequence variants amplified to a discernible extent by the depth of Illumina sequencing. But the distinctiveness of some of these OTUs from obviously host-derived sequences was so great in some cases (e.g., the strongly *Fucus*-associated ‘Uncharacterised’ clade in Fig. 7) that we propose that these represented hitherto unknown epi-endophytes. In other cases we have conservatively ascribed OTU diversity clustering with host sequences (e.g., the clades bracketed as *A. nodosum* and *H. elongata* in Fig. 7) as being host-derived, but there remains the possibility that they represented different organisms.

Some of the brown algal OTUs detected in our study, such as the ones clustering with the epiphytic *Pylaiella* and *Halothrix* and other brown algal epi-endophytes (Fig. 7), also represented epi-endophytic lineages [e.g., 48, 58, 81]. These OTUs were detected in all the brown algal hosts included in this study, indicating a broad distribution and host range.

By analysing sequences from cultured phaeophyte symbionts of brown algae and red algae (Attwood 2005, [141]) we show that the epi-endophyte genera *Microspongium*, *Myriotrichia*, *Myrionema*, and *Streblonema* form a clade with *Chorda* and *Ectocarpus* (growing as epi-endophytes), and previously sequenced brown algal epi-endophytes *Pylaiella*, *Halothrix*, and *Scytosiphon*, which all together form a maximally supported clade with *Saccorhiza*. This, however does not account for the vast majority of host-associated brown algal sequences in our data that are not obviously host-derived. Further culturing and *is situ* experiments are required to explain our findings. One possible explanation is that these brown algal sequences might represent microscopic life-stages (zoospores, gametophytes, gametes and juvenile sporophytes) of brown algae growing epi-/endophytically on substrata, including on macroscopic sporophytes of other seaweeds [e.g., 82, 83-85]. This can also explain why the host-derived OTUs were detected in lower abundance in the other brown algal samples (Fig. 7).

Another possible explanation is that small filamentous brown algae were growing epi-endophytically in the larger brown algal hosts [48, 86]. Such endophytic infections are described as common diseases of brown algae [e.g., 52, 53, 67, 68, 87]. Hosts containing brown algal endophytes can either be asymptomatic, show weak symptom such as dark spots or warts, or present severe symptoms such as morphological changes to the brown algal host tissues [52, 67]. Substrata and brown algal thalli often contain many more brown algal endophytes than expected based on visual inspections prior to culturing [47, 53, 86]. The majority of these previous studies have, however, either not used molecular methods or have used different marker regions than 18S, and consequently there are no reference sequences available in databases to compare with our sequence data.

The green algal OTUs were all similar to reference sequences of taxa that are known to be endophytes in red (*Chondrus crispus Mastocarpus* stellatus, *Osmundea* spp. and *Dumontia contorta*) and brown algae (*Fucus serratus* and *Chorda filum*) [33, 88, 89].

All red algal OTUs were similar to reference sequences of taxa that are common epiphytes on seaweeds and a variety of other substrata. Several of the taxonomic names for the reference sequences included in the phylogeny are out of date (Fig. 7): *Ceramium* spp. is an aggregate of species (i.e. *Ceramium rubrum* in Fig. 7) which are members of the order Ceramiales together with *Calithamnion* spp., *Polysiphonia* sensu lato (*Polysiphonia fucoides* is now *Vertebrata fucoides*) and *Cryptoleura* spp. Red algal endophytes commonly infect other, larger red algae [e.g., 90]. We did not, however find any endophytic red algae in the brown algae investigated in our study. This might indicate that brown algae are not prone to endophytic infections by red algae, but instead are more susceptible to endophyte infestations by other brown algae.

### Novel diversity and putative parasites

We demonstrate that brown algal holobionts encompass potential novel eukaryotic diversity, as almost one third of the OTUs we detected have not been found in molecular environmental surveys before. This was more pronounced in some phylogenetic groups than in others, in particular the labyrinthulids and oomycetes (Fig. 3), Cercozoa (Fig. 4), Amoebozoa and Centroheliozoa (Supplementary Fig. 3). Some of the OTUs were taxonomically and phylogenetically placed with known and putative parasites of seaweeds. Our findings and the fact that the vast majority of macroalgae in marine habitats remain unexplored, demonstrates that brown algae and other seaweeds are potentially rich sources for a large and hidden diversity of novel microeukaryotes.

Labyrinthulomycetes are heterotrophic stramenopiles which are abundant and diverse in a wide range of marine and freshwater habitats where they play important roles as saprotrophs/decomposers [91, 92]. Some Labyrinthulomycetes are also known as important symbionts (parasites, mutualists or commensals) of marine organisms, including brown algal seaweeds [93–95]. Although most thraustochytrids are free living, a few species have been associated with disease in marine metazoans, including the quahog parasite QPX which parasitizes clams. Several of the OTUs in our study clustered in three highly supported clades within Labyrinthulida (Fig 3), and had high similarity to *Aplanochytrium* labyrinthulids associated with various hosts such as corals [96], the pseudoparenchymatous brown alga *Elachista* sp. [95], seagrass [97] and sea stars [98].

Certain oomycetes are common parasites of brown and red algae [37, 99–101], and *Olpidiopsis* species have been shown to have cosmopolitan occurrence and broad host ranges [100, 102]. In this study, we found OTUs clustering within Olpidiopsidales. Some of these were highly similar to reference sequences of *Anisolpidium rosenvingei* and *A*. *ectocarpii* (Fig 3), which are parasites of filamentous brown algae [36]. Others displayed high similarity to *Pontisma lagenidioides* and “*Ectrogella*” (lnfLicSC1, SC2-a, SC3 and SC4) which parasitizes the diatom *Licmophora* sp. and the red algae *Ceramium virgatum* [103, 104]. Further, we detected two novel OTUs with no close relatives among available reference sequences (OTU 214; 90.9 % identity to JN635125 and OTU 290; 92.4 % identity to KT273921, Fig. 3), of which the closest known relative to the latter is an oomycete parasitizing diatoms [103, 105].

Cercozoan diversity was dominated by vampyrellid amoebae, with four OTUs forming a distinct clade with GU385680, which was previously detected on *Ascophyllum nodosum* used as live bait wrapping [80]. Vampyrellids exhibit a wide diversity of feeding strategies and often feed omnivorously; it is striking that this clade comprises only lineages associated with brown algae, suggesting a specific association. Phytomyxids, which are biotrophic parasites of angiosperms, oomycetes, and stramenopile algae [39], were represented by two OTUs in a clade of marine sediment-derived sequences. Consequently, there is no direct evidence for that these phytomyxids have a specific interaction with macroalgae. OTU 242 grouped with endomyxan Novel Clade 9 [106], members of which are found in a wide range of environments and have been hypothesized to be parasites of algae.

Fungi associated with brown algae can have everything from detrimental to beneficial effects on their hosts [107, and references therein]. Two of the Ascomycota OTUs clustered with taxa known to be parasites of brown algae such as *Sarocladium* and *Acremonium* [108] (Fig. 5). Several of the basidiomycete OTUs clustered with reference sequences of *Cryptococcus* and *Cystofilobasidium* that have previously been found associated to various marine invertebrates, algae and seaweeds (Fig. 5) [109–112]. In addition, within Chytridiomycota there were several OTUs that clustered with *Rhizophydium* sp., which comprise parasites with broad host ranges known to infect both macroalgae and protists [113, 114]. One OTU (OTU 76; Fig. 5) also displayed high similarity to *Chytridium polysiphoniae*, a parasite of brown algae [115, 116]. The final chytrid OTU clustered with *Mesochytrium penetrans*, a parasite of the unicellular green algae *Chlorococcum minutum* [117], indicating that this could represent a parasite of one of the green algae found in our study, or that relatives of *M*. *penetrans* have a broader host range than currently known.

## Conclusions

Brown algal holobionts are major marine habitat formers that provide food and shelter for diverse communities of both macro- and microorganisms. Our findings demonstrate that microeukaryotes should be considered as an integral part of brown algal holobionts, and provides important baseline data for future studies of seaweed-associated microorganisms. The potential novel eukaryotic diversity we found and the fact that the vast majority of macroalgae in marine habitats remain unexplored, demonstrates that brown algae and other seaweeds are potentially rich sources for a large and hidden diversity of novel microeukaryotes and epi-endophytes.

The prerequisite to understand the ecology of brown algae and their host-associated microbes, is to know their diversity, distribution and community dynamics. Consequently, in order to obtain an understanding of the interdependence of all the different players in brown algal holobionts it is critical to include the so far largely neglected eukaryotic and microeukaryotic partners in future holobiont investigation. It is however also important to remember that microeukaryotes interact not only with the host but also with the prokaryotes and viruses that share the same host environment, and future holobiont surveys should ideally include all of these components.

## Methods

### Sample collection

Five individuals each of *Ascophyllum nodosum*, *Fucus vesiculosus*, *Laminaria digitata* and *Saccharina latissima* were sampled from the Oslofjord (59°40′26.9″N, 10°35′13.199″E), while five individuals each of *F*. *vesiculosus*, *F*. *serratus Himanthalia elongata*, *Saccorhiza polyschides* and *L*. *digitata* were collected at Newton’s Cove (Dorset, UK: 50°36′18″N, 2°26′58″W) in October 2015. Samples were collected by free diving and shore-based collection, kept cool in separate containers of seawater while transported to molecular laboratories where they were immediately subjected to subsampling of tissues for molecular analyses.

All samples were handled under laminar flow hoods to limit their exposure to airborne contaminants. The brown algae were rinsed in sterile artificial seawater (ASW) to remove loosely attached organisms (but not true epibionts) and debris from the surface. This was done in three consecutive steps by vortexing the samples in sterile 50 mL tubes with ASW for 5-10 seconds. The rinsed individuals were placed in sterile petri dishes, and one subsample of the algal thallus (approximately 1 cm^2^ squares) per individual was excised and placed in separate 2 mL tubes. All algae looked healthy; we did not specifically target potentially infected tissues.

The samples were freeze-dried under sterile conditions by perforating the sample tube lids and placing the sample tubes inside sterile culture flasks with 0.2 um filter caps (Thermo Scientific^TM^ Nunclon^TM^). After freeze-drying, samples were weighted, placed in new 2 mL sample tubes, and stored at −80 °C until DNA extraction.

### Endophyte sampling

As part of a previous student project (Attwood 2005, [141]), brown algal endophytes were selected from the B. Rinkel collection at The Natural History Museum, London. The samples were isolated from hosts as shown in Table 5. Partial 18S-ITS1 rDNA sequences were amplified using the phaeophyte-specific primer DICI [118] and the general primer TW7 [119]. The 18S region of this amplicon did not overlap with the V4 region sequenced by the metabarcoding approach, so longer 18S amplicons were generated from the brown macroalgal samples used for this study as described below, to enable as much direct comparison of our newly generated sequences with those from the endophytes. Full details of the DNA extraction, PCR, and sequencing methods used for the endophytes are given in Attwood 2005 [141].

**Table 5.** Hosts with brown algal endophytes selected for isolation from the B. Rinkel collection at the Natural History Museum, London.

| Sample ID | Host | Site name, county | Date | Collector |
| --- | --- | --- | --- | --- |
| BRE1sn1 | *Chondrus crispus* | Sheerness, Kent | 22.10.03 | J. Brodie |
| BRE9ls2 | *Chondrus crispus* | Lilstock, Somerset | 27.10.03 | B. Rinkel |
| BRE38sm11 | *Mastocarpus stellatus* | Sidmouth, Devon | 28.10.01 | B. Rinkel |
| BRE74ls1 | *Fucus serratus* | Lilstock, Somerset | 27.10.03 | B. Rinkel |
| BRE98sm12 | *Mastocarpus stellatus* | Sidmouth, Devon | 28.10.01 | B. Rinkel |
| BRE117sm22 | *Osmundea osmunda* | Sidmouth, Devon | 28.10.01 | B. Rinkel |
| BRE118sm22 | *Osmundea osmunda* | Sidmouth, Devon | 28.10.01 | B. Rinkel |
| BRE128hi1 | *Osmundea pinnatifida* | Hayling Island, Hampshire | 05.02.04 | B. Rinkel |
| BRE152cb2 | *Chondrus crispus* | Castle Beach, Pembrokeshire | 09.03.04 | B. Rinkel |
| BRE157br1 | *Mastocarpus stellatus* | Black Rock, Pembrokeshire | 10.03.04 | B. Rinkel |
| E026pz | *Chondrus crispus* | Polzeath, Cornwall | 19.06.04 | M. Holmes |
| E030Nb | *Mastocarpus stellatus* | Norway | 23.06.04 | J. Brodie |
| E326wr | *Fucus serratus* | West Runton, Norfolk | 28.09.04 | B. Rinkel |
| E360sm2 | *Chorda filum* | Sidmouth, Devon | 28.10.03 | B. Rinkel |
| E361ac | *Chorda filum* | A'Chleit, Bute | 10.07.04 | B. Rinkel |
| E365sg | *Mastocarpus stellatus* | Seagreens, Aberdeenshire | 16.07.04 | B. Rinkel |
| E368 | *Chondrus crispus* | Bouley Bay, Jersey | 13.03.05 | B. Rinkel |
| E370 | *Chondrus crispus* | Bouley Bay, Jersey | 13.03.05 | B. Rinkel |
| E377 | *Chondrus crispus* | Greve de Lacq, Jersey | 14.03.05 | B. Rinkel |
| LTE19/1 | *Osmundea pinnatifida* | Black Rock, Pembrokeshire | 10.03.04 | B. Rinkel |

### Molecular methods

The freeze-dried samples were mechanically disrupted by adding two sterile tungsten carbide beads to each sample tube and tissue-lyzed at 20Hz for 2 minutes, or longer, until the tissues were completely pulverised. Thereafter, a modified lysis buffer [120] was added to the tubes proportional to sample weight. This lysis buffer contains antioxidant compounds such as PVP (Polyvinylpyrrolidone) and BSA (Bovine Serum Albumin) that bind polyphenols, and has high salt concentration which decrease the levels of coextracted polysaccharides. The samples were homogenised and divided in two or more tubes so that the sample used for DNA isolation did not exceed 20 mg dry weight tissue. DNA was extracted following the protocol by Snirc *et al*. [120].

PCR amplification of the V4 region of the 18S rDNA gene was performed with ‘pan eukaryotic’ primers; the forward primer used was a slightly modified version of the TAReuk454FWD1 primer (5’-CCAGCASCYGCGGTAAT**<u>W</u>**CC-3’) and reverse primer TAReukREV3 (5’-ACTTTCGTTCTTGATYRA-3’) [121]. PCR reactions (25 μl) contained 1x KAPA HiFi HotStart ReadyMix (Kapa Biosystems), 0.4 μM of each primer, 0.5 mg/mL Bovine serum albumin (BSA; Promega) and 1 μl genomic DNA. The PCR program had an initial denaturation step at 98°C for 2 min, 15 cycles of 30 s at 98°C, 30 s at 53°C and 45 s at 72°C, then 20 similar cycles except that the annealing temperature was 48°C, and a final elongation step at 72°C for 10 min. DNA was titrated into several dilutions to take into account potential inhibiting substances from the brown algae. Each brown algal individual sample was amplified separately. PCR products were purified and eluted (20 μl) with ChargeSwitch PCR Clean-Up kit (ThermoFisher Scientific), and quantified with the dsDNA BR Assay Kit and Qubit 2.0 Fluorometer (ThermoFisher Scientific), before they were pooled equimolarly, according to species and geographic location; e.g., the purified PCR products of five *Fucus vesiculosus* individuals from Norway were pooled into one sample before sequencing library preparation.

Sequencing libraries were prepared using the TruSeq Nano DNA Library Preparation Kit at The Natural History Museum, London UK, following the manufacturer’s protocol. High throughput sequencing was conducted on a MiSeq v3 flow cell using 2×300 bp paired-end reads. Base calling was performed by Illumina Real Time Analysis v1.18.54 and was demultiplexed and converted to FastQ files with Illumina Bcl2fastq v1.8.4.

The complete sequencing dataset is available at the European Nucleotide Archive under the study accession number XXXXXXXX (http://www.ebi.ac.uk/ena/ data/view/ XXXXXXXX).

To obtain full length 18S sequences of the brown algal host species, three DNA isolates from each brown algal species were subjected to PCR amplification using the general eukaryotic 18S primers NSF83 (5’-GAAACTGCGAATGGCTCATT-3’ [122]) and 1528R (5’-TCCTTCTGCAGGTTCACCTAC-3’ [123]) utilising illustra™ PuReTaq Ready-To-Go™ PCR beads (GE Healthcare). PCR reactions contained 1 Illustra PuReTaq Ready-To-Go-bead, 1μl of DNA template, 0.5 μM of each primer, filled to 25μl with water. PCR settings were: initial denaturation at 94°C for 2 min; 35 cycles of denaturation at 94°C for 45 s, annealing at 55°C for 30 s, elongation at 68°C for 1.5 min; final elongation step at 68°C for 15 min. The PCR products were cloned using the TOPO TA Cloning Kit for Sequencing (Invitrogen), according to the manufacturer’s protocol. The clones were grown overnight in Luberia-Bertoni (LB) media amended with 50 μg/mL ampicillin. From each brown algal species, 20-30 bacterial colonies/cloned fragments, were subjected to PCR reactions with the vector primers T7 and M13R and using ca 0.5 μl of the bacterial suspension as template. PCR settings were: initial denaturation at 94°C for 10 min; 30 cycles of denaturation at 94°C for 30 s, annealing at 53°C for 1 min, elongation at 72°C for 2 min; final elongation step at 72°C for 10 min. The PCR products were cleaned using illustra™ ExoProStar 1-Step (GE Healthcare) and then Sanger sequenced (GATC Eurofins Genomics, Germany).

### Bioinformatics

The sequence data was processed using a workflow for MiSeq amplicon data (https://github.com/ramalok/amplicon_processing). In short, prior to downstream analyses error correction was done with BayesHammer [124], before merging the paired-end reads with PEAR [125]. Thereafter, quality filtering, length check and fasta conversion (from fastq to fasta) was done with USEARCH [126] before reads were put into the correct 5’ – 3’ direction using profile hidden Markov models (hmmer-3.0rc2). Dereplication, sorting by abundance and discarding of singletons was done using USEARCH 64bit version. Clustering of reads into Operational Taxonomic Units (OTUs) was done using the UPARSE algorithm as implemented in USEARCH, applying a 97% sequence similarity threshold. *De-novo* and reference-based chimera checking was performed and chimeric sequences discarded. OTUs were taxonomically assigned using BLASTn against the PR2 v. 4.12.0 [127], GenBank [128] and SILVA SSU [129] databases. We used a relaxed BLAST search strategy (BLASTn parameters: -evalue 0.0001 -perc_identity 75) to retrieve distant sequences, and only retrieved the best hit per query sequence. The PR2 taxonomy was used for taxonomy assignment for the majority of the following analyses.

### “Post-UPARSE trimming” & Diversity analyses

Species (OTU) composition analyses and further trimming of the dataset was done using R v. 3.6.3 (R Core Team 2013). As we expected the sequence dataset was heavily dominated by brown algal host reads, and we did therefore not normalise the dataset because this would have led to removal of true diversity since most of the other OTUs had very low read numbers compared to the brown algal OTUs. Further, all OTUs assigned to Metazoa were removed from the dataset in addition to OTUs assigned to land plants. The 57 OTUs (16.3% of the total number of OTUs) that matched those of Metazoa are shown in Supplementary Figure S5, Supplementary Table S5. The trimmed dataset, containing only non-metazoan eukaryotes, was used for further analyses.

The taxonomy of OTUs was visualized using the ggplot2 package v. 3.3.2. [130]. To investigate the composition of (micro)eukaryotes associated with the different brown algal hosts, Bray-Curtis pairwise distances were calculated using vegan v. 2.5-6 [131] for all brown algal samples (including/excluding brown algal OTUs) and analysed using non-metric multidimensional scaling (NMDS) (ggplot2 package v. 3.3.2 [130]). Analysis of similarities [ANOSIM; 132] was calculated using vegan v. 2.5-6 [131] to test if there were statistically significant differences between clusters of libraries corresponding to different brown algal host genera. The similarity percentage [SIMPER; 132] was calculated using the simper.pretty.R script [133] to identify OTUs that contributed the most to the dissimilarities observed. Indicator species analyses (IndVal) were carried out using the indicspecies package v. 1.7.9. [134], to identify OTUs preferentially associated with specific brown algal host genera. Venn diagrams intersects were computed using the function overLapper and plotted using the function vennPlot in the R-package systemPipeR v. 1.24.3 [135].

### Phylogenetic analysis

All OTUs were places into a global eukaryotic alignment of near full-length 18S reference sequence using MAFFT v7.300b with the ‘einsi’ option for global homology and long gaps for the reference sequences [136]. The shorter OTUs from the V4 region (300-450 bp) were added to the reference alignment using the –addfragment option in MAFFT and ambiguously aligned sites were removed with TrimAl [137] with –gt 0.3 and –st 0.001. Phylogenetic trees were built with RAxML v8.0.26 under the GTRGAMMA model and the autoMRE bootsopping criteria [138]. Based on a combination of the phylogenetic affinities of the OTUs and the taxonomic assignment from PR2, individual phylogenies of the most abundant/diverse groups of microeukaryotes – and those with known associations with brown algae - were constructed: i.e Fungi, oomycetes & labyrinthulids, Cercozoa, diatoms, red, brown, and green algae, ciliates, other alveolates, Amoebozoa, and centroheliozoans.

Individual sequence datasets were constructed including OTUs from each group whose taxonomic annotation and phylogenetic position were concordant, and their closest Blastn matches from GenBank. Sequence alignments were carried out in MAFFT as described above. Phylogenetic trees were built with MrBayes v.3.2.6 [139]. Two separate MC3 runs with randomly generated starting trees were carried out for 4 million generations each with 1 cold and 3 heated chains. The evolutionary model applied a GTR substitution matrix, with a 4-category autocorrelated gamma correction. All parameters were estimated from the data. The trees were sampled every 1000 generations and the first 1 million generations discarded as burn-in. All phylogenetic analyses were carried out on the Cipres server [140]. Heatmaps were made for all phylogenetic trees to display the proportional read abundance (log_10_) for each OTU in the different samples.

## Declarations

### Ethics approval and consent to participate

Not applicable

### Consent for publication

Not applicable

### Availability of data and materials

The dataset generated are available in XX repository, under the accession XXXX-XXXX (link to dataset).

### Authors’ contributions

**MFMB**, **DB** & **KST** planned and designed the study. **MFMB**, **JF**, **SF** & **AWD** planned, organised sampling and collected samples. **DB** collected samples from the UK. **MFMB**, **JF** & **AWD** processed all samples (molecular lab work). **JB** provided the phaeophyte endophyte data generated by **SA**. **RL** did the initial bioinformatic processing of the raw sequence dataset. **MFMB**, **DB** & **AKK** analysed the data. **MFMB**, **DB** & **JB** interpreted the results and wrote the manuscript. All authors contributed substantially to manuscript revisions. All authors read and approved the final manuscript.

## Acknowledgements

We thank Jonas Thormar, Jens Ådne Haga and Ane-Rikke N. Oen-Ting for help with field sampling of brown algae in Oslofjord. Drøbak Biological Station and Hans Erik Karlsen are thanked for providing sampling equipment and laboratory facilities for sample preparations.

## Supplementary figures

**Supplementary figure 1. Ciliates: phylogeny with heatmap representing proportional read abundance (log_10_) of OTUs per sample**. Bayesian phylogeny of Ciliates with bootstrap values from maximum likelihood analysis added. Black circles represent strong support (posterior probability >/= 0.95, and bootstrap support >/= 95%). OTUs from this study are shown in bold. The percentage identity to the most similar reference sequence is shown in parenthesis after each OTU. Host or type of environment the reference sequences were retrieved from in previous studies are listed in parenthesis after each GenBank accession number. The scale bar represents 0.2 substitutions per site. Abbreviations used for descriptions of environment: env.S = environmental sample, marine sediment; env.W = environmental sample, marine water; env.FW = environmental sample, fresh water; env.WW = environmental sample, waste water; env.BW = environmental sample, brackish water; env.S/W = environmental sample, marine sediment or water; env.MBF = environmental sample, marine biofilm; env.FBF = environmental sample, freshwater biofilm; env.So = environmental sample, soil. The heatmap illustrates the log_10_ read abundance for each OTU in the brown algal samples of *Fucus serratus*, *F*. *vesiculosus*, *Ascophyllum nodosum*, *Himanthalia elongata*, *Saccorhiza polyschides*, *Laminaria digitata* and *Saccharina latissima*. The sampling location for each brown alga is shown in parenthesis; NO = Norway and UK = The United Kingdom. All samples were collected in October 2015, except *F*. *vesiculosus* (NO*) which was sampled in Norway, May 2013.

**Supplementary figure 2. Apicomplexa: phylogeny with heatmap representing proportional read abundance (log_10_) of OTUs per sample**. Bayesian phylogeny of Apicomplexa and perkinsids with bootstrap values from maximum likelihood analysis added. Black circles represent strong support (posterior probability >/= 0.95, and bootstrap support >/= 95%). OTUs from this study are shown in bold. The percentage identity to the most similar reference sequence is shown in parenthesis after each OTU. Host or type of environment the reference sequences were retrieved from in previous studies are listed in parenthesis after each GenBank accession number. The scale bar represents 0.3 substitutions per site. Abbreviations used for descriptions of environment: env.S = environmental sample, marine sediment; env.W = environmental sample, marine water; env.FW = environmental sample, fresh water; env.I = environmental sample, ice; GC = gut content. The heatmap illustrates the log_10_ read abundance for each OTU in the brown algal samples of *Fucus serratus*, *F*. *vesiculosus*, *Ascophyllum nodosum*, *Himanthalia elongata*, *Saccorhiza polyschides*, *Laminaria digitata* and *Saccharina latissima*. The sampling location for each brown alga is shown in parenthesis; NO = Norway and UK = The United Kingdom. All samples were collected in October 2015, except *F*. *vesiculosus* (NO*) which was sampled in Norway, May 2013.

**Supplementary figure 3. Amoebozoa and Centroheliozoa phylogeny with heatmap representing proportional read abundance (log_10_) of OTUs per sample**. Bayesian phylogeny of Amoebozoa and Centroheliozoa with bootstrap values from maximum likelihood analysis added. Black circles represent strong support (posterior probability >/= 0.95, and bootstrap support >/= 95%). OTUs from this study are shown in bold. The percentage identity to the most similar reference sequence is shown in parenthesis after each OTU. Host or type of environment the reference sequences were retrieved from in previous studies are listed in parenthesis after each GenBank accession number. The scale bar represents 0.2 and 0.08 substitutions per site for Amoebozoa and Centroheliozoa respectively. Abbreviations used for descriptions of environment: env.FWS = environmental sample, freshwater sediment; env.W = environmental sample, marine water; env.So = environmental sample, soil. The heatmap illustrates the log_10_ read abundance for each OTU in the brown algal samples of *Fucus serratus*, *F*. *vesiculosus*, *Ascophyllum nodosum*, *Himanthalia elongata*, *Saccorhiza polyschides*, *Laminaria digitata* and *Saccharina latissima*. The sampling location for each brown alga is shown in parenthesis; NO = Norway and UK = The United Kingdom. All samples were collected in October 2015, except *F*. *vesiculosus* (NO*) which was sampled in Norway, May 2013.

**Supplementary figure 4. Dinoflagellates: phylogeny with heatmap representing proportional read abundance (log_10_) of OTUs per sample**. Bayesian phylogeny of Dinoflagellates with bootstrap values from maximum likelihood analysis added. Black circles represent strong support (posterior probability >/= 0.95, and bootstrap support >/= 95%). OTUs from this study are shown in bold. The percentage identity to the most similar reference sequence is shown in parenthesis after each OTU. Host or type of environment the reference sequences were retrieved from in previous studies are listed in parenthesis after each GenBank accession number. The scale bar represents 0.08 substitutions per site. Abbreviations used for descriptions of environment: env.S = environmental sample, marine sediment; env.W = environmental sample, marine water; env.BW = environmental sample, brackish water. The heatmap illustrates the log_10_ read abundance for each OTU in the brown algal samples of *Fucus serratus*, *F*. *vesiculosus*, *Ascophyllum nodosum*, *Himanthalia elongata*, *Saccorhiza polyschides*, *Laminaria digitata* and *Saccharina latissima*. The sampling location for each brown alga is shown in parenthesis; NO = Norway and UK = The United Kingdom. All samples were collected in October 2015, except *F*. *vesiculosus* (NO*) which was sampled in Norway, May 2013.

**Supplementary figure 5. Diversity of metazoan OTUs detected in brown algal holobionts.** Proportional abundance of OTUs taxonomically assigned to Metazoa in the brown algal samples of *Fucus serratus*, *F*. *vesiculosus*, *Ascophyllum nodosum*, *Himanthalia elongata*, *Saccorhiza polyschides*, *Laminaria digitata* and *Saccharina latissima*. The sampling location for each brown alga is shown in parenthesis; NO = Norway and UK = The United Kingdom. The asterisk (NO*) represent *F*. *vesiculosus* sampled in Norway, May 2013. All other samples were collected in October 2015.

