## Supplementary figures and images for "Elucidating the diversity of microeukaryotes and epi-endophytes in the brown algal holobiome"

### Supplementary Fig. 1

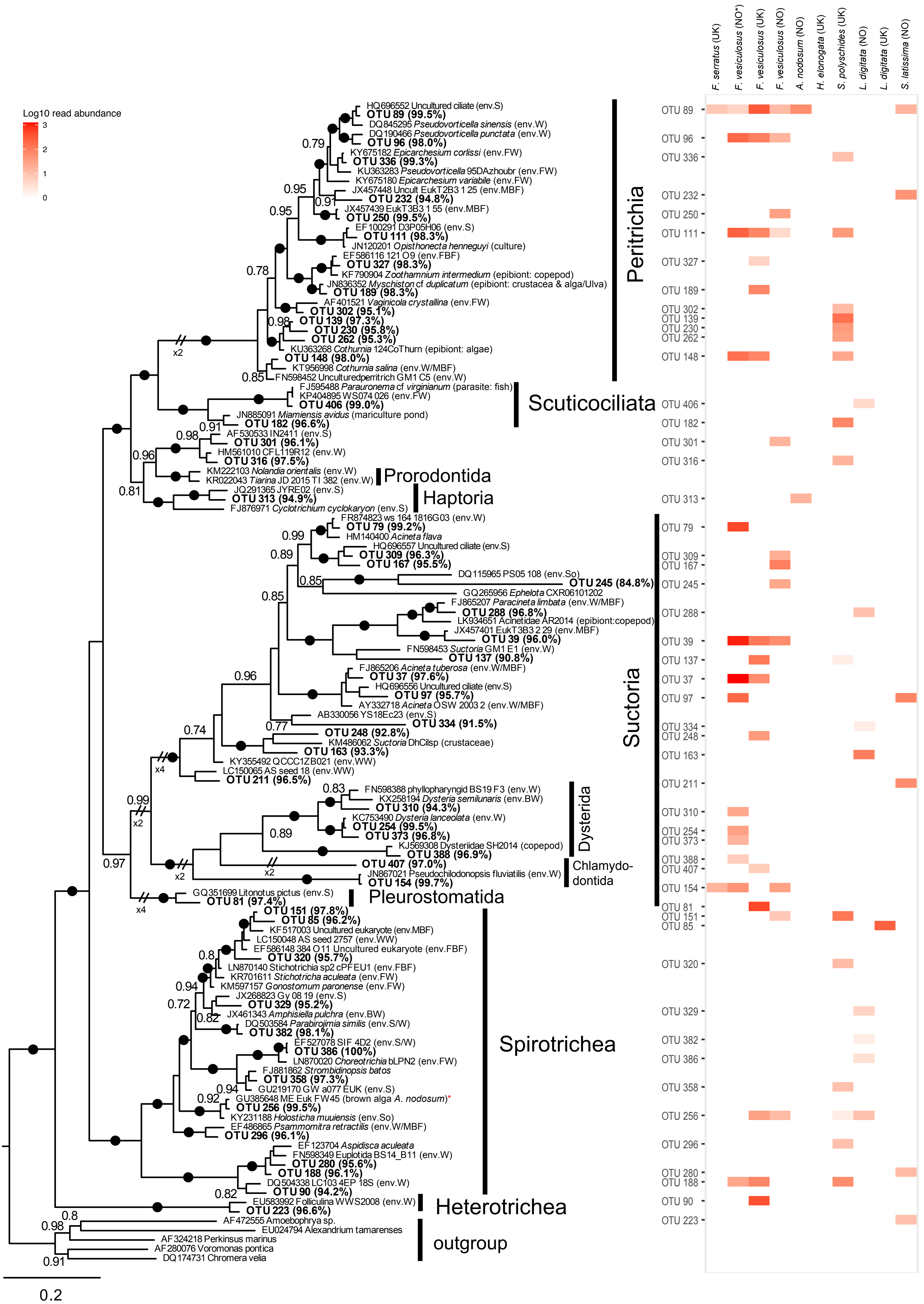

### Supplementary Fig. 2

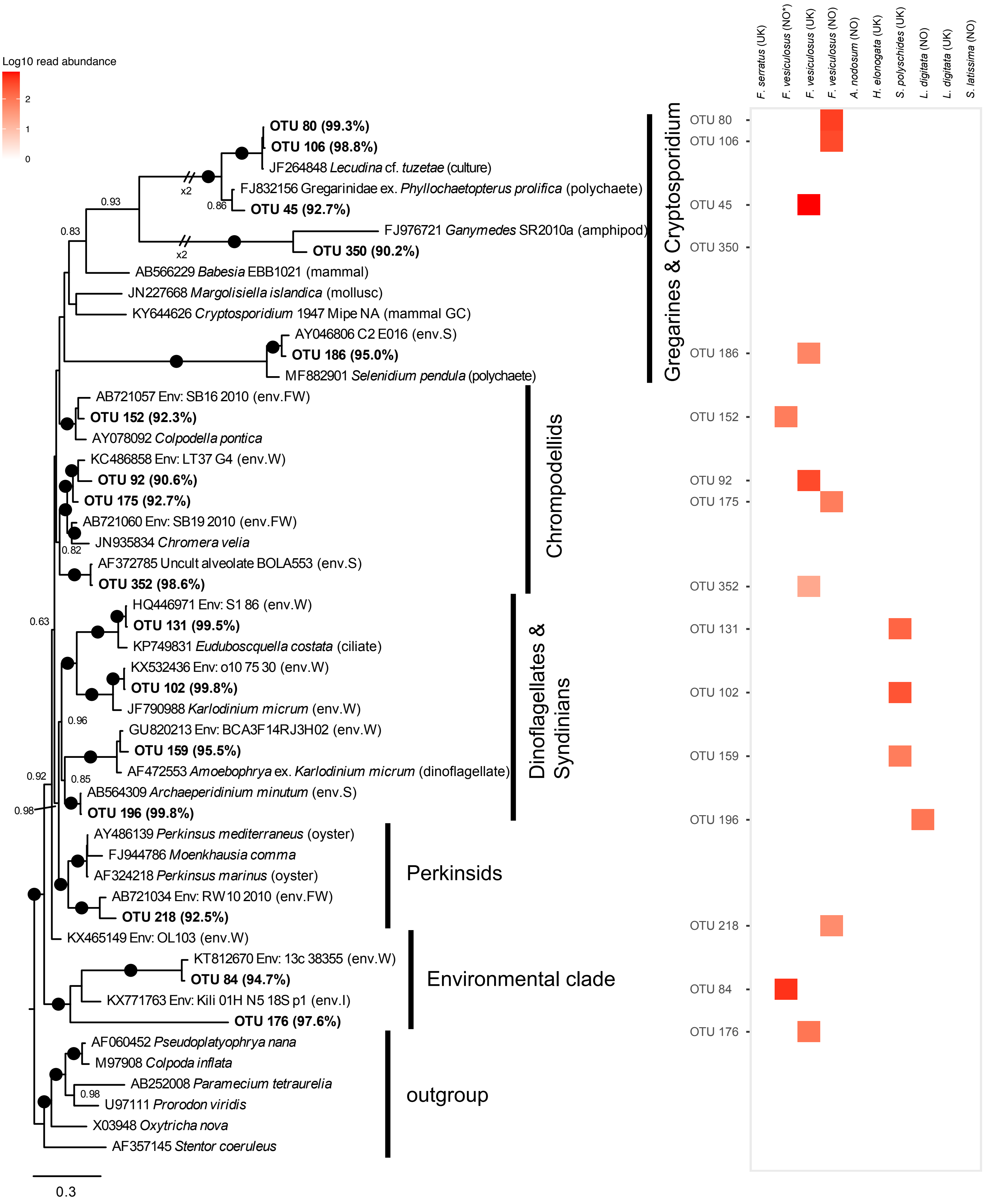

### Supplementary Fig. 3

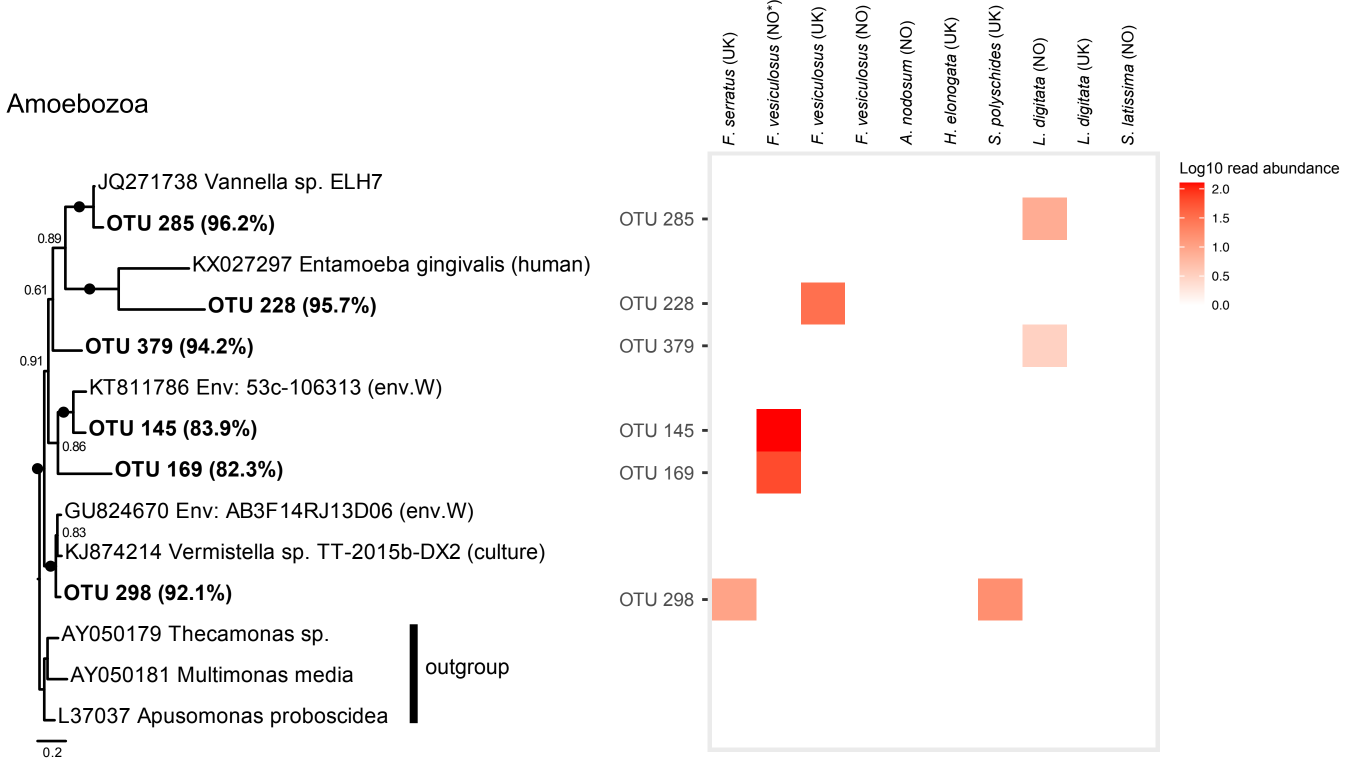

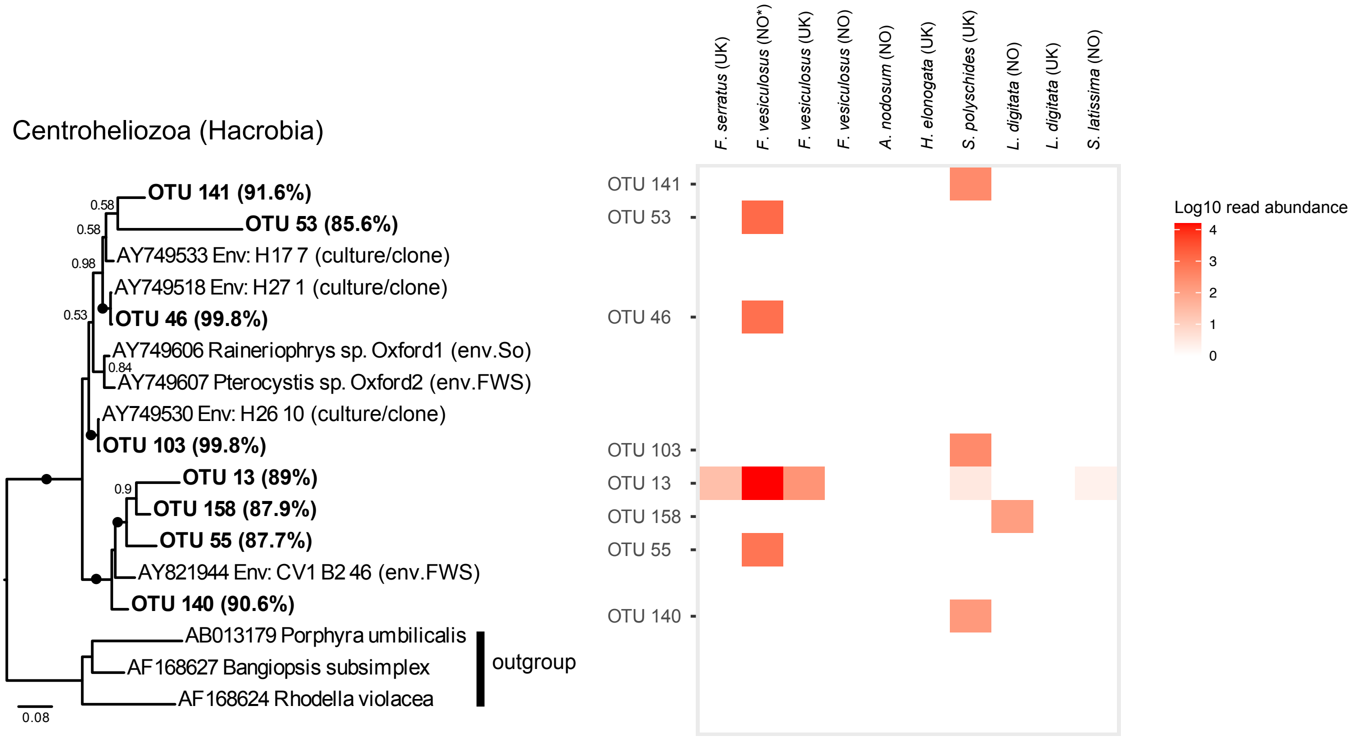

### Supplementary Fig. 4

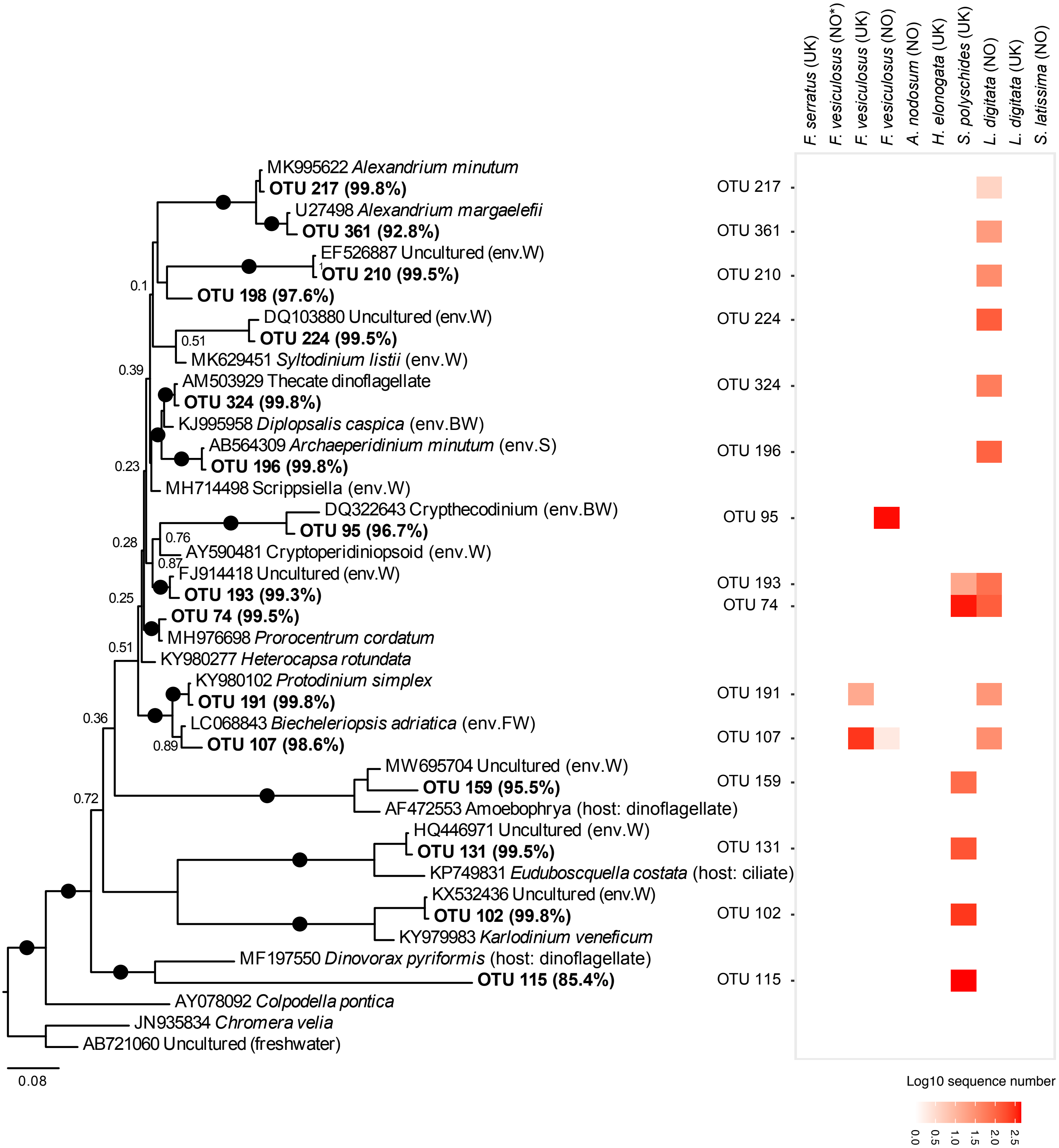

### Supplementary Fig. 5

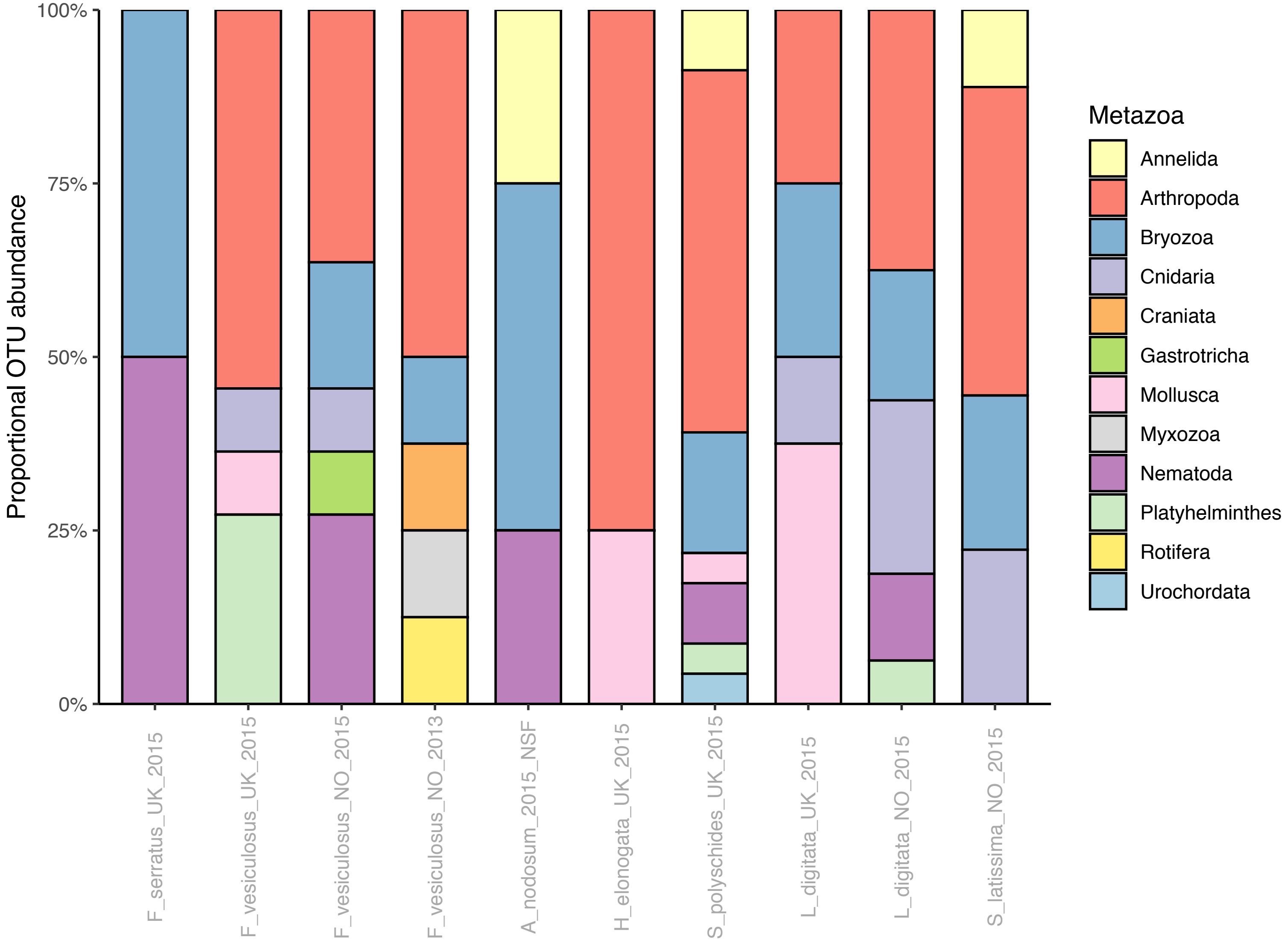
